# CD133-dependent membrane organization shapes extracellular vesicle heterogeneity in triple-negative breast cancer

**DOI:** 10.64898/2026.11.20.768123

**Authors:** Mireia Gomez-Duro, Ptisssam Bergam, Anne-Sophie Macé, Clotilde Théry, Lorena Martin-Jaular, Gisela D’Angelo

## Abstract

Extracellular vesicle (EV) heterogeneity is well established, but how distinct EV populations emerge from the same donor cell remains poorly understood. Here, we investigated plasma membrane organization as a potential contributor to EV molecular heterogeneity in triple-negative breast cancer (TNBC). CD133/PROM1 expression was associated with plasma membrane remodeling, including prominent protrusion formation, and with the redistribution of the EV-associated tetraspanin CD63 toward these domains. Quantitative immunogold electron microscopy revealed a disproportionate enrichment of CD63 labeling within protrusions following CD133 expression. This remodeling was accompanied by increased EV particle output and altered EV-associated marker signals. Single-particle analysis identified distinct CD133 /CD63, CD133 /CD63, and CD133 /CD63 EV populations, demonstrating that CD133 and CD63 were not uniformly distributed across detected EVs. CD63 was detected more frequently in the CD133 than in the CD133 EV fraction, supporting an association between CD133 expression and EV marker heterogeneity. CD133 expression was also associated with changes in intracellular multivesicular body (MVB) density and size distribution, with different responses across the two TNBC cell models examined. Together, these findings suggest that plasma membrane organization may contribute to EV heterogeneity alongside context-dependent remodeling of intracellular MVB compartments. Rather than implicating CD133 in a single EV biogenesis pathway, our data highlight spatial membrane organization as a potential determinant of EV molecular diversity.

## INTRODUCTION

Extracellular vesicles (EVs) are membrane-enclosed particles released by virtually all cell types and important mediators of intercellular communication. By transferring proteins, lipids and nucleic acids between cells, EVs can modify the phenotype and behavior of recipient cells and contribute to physiological and pathological processes, including tumor progression. EVs comprise a heterogeneous group of vesicles that differ in their molecular composition, biophysical properties and cellular origin (van Niel et al., 2018; van de Wakker et al., 2023; Carney et al., 2025; D’Angelo et al., 2025). Their biogenesis can involve distinct membrane trafficking pathways, including outward budding and fission of the plasma membrane and inward budding of endosomal membranes followed by the formation and secretion of intraluminal vesicles (ILVs) from multivesicular bodies (MVBs). These pathways are not necessarily mutually exclusive and can generate EV populations with distinct molecular compositions and biological properties (Dixson et al., 2023; Welsh et al., 2024).

The cellular mechanisms controlling the relative contribution of these pathways remain incompletely understood. In particular, the organization of the plasma membrane and the spatial distribution of membrane proteins and lipids may influence where vesicle formation occurs and which molecular components are incorporated into released EVs (Dixson et al., 2023). The plasma membrane is a highly organized and dynamic structure in which specific lipids and proteins are concentrated within distinct membrane domains that regulate signaling, cytoskeletal organization and membrane trafficking (Dixson et al., 2023). Membrane curvature and cortical cytoskeletal organization are particularly prominent features of membrane protrusions, including microvilli- and filopodia-like structures. Increasing evidence indicates that such protrusive domains can be associated with EV shedding, suggesting that plasma membrane architecture may contribute to the generation of molecularly distinct EV populations (Rilla 2021; Nishimura et al. 2021; D’Angelo et al. 2023; Gomez-Duro et al., 2025; Fujioka et al., 2025).

Protrusion-associated EV release has been described in several cellular contexts, and the enrichment of specific proteins and lipids within protrusive membrane domains suggests that these structures may provide spatially organized sites for vesicle formation and cargo selection (Marzesco et al., 2009; Corbeil et al., 2010; D’Angelo et al., 2023; Fujioka et al., 2025). Membrane curvature, actin organization and the local concentration of membrane proteins may contribute to the formation and release of vesicles from these domains (D’Angelo et al., 2023). However, the molecular mechanisms that establish these membrane domains and determine the selective recruitment of EV-associated proteins remain poorly understood. In particular, whether the spatial organization of the donor-cell plasma membrane contributes to the molecular heterogeneity of released EVs remains an open question.

Prominin-1 (CD133/PROM1) is a pentaspan transmembrane glycoprotein enriched in specific plasma membrane protrusions and associated with cholesterol-rich membrane microdomains. Its localization to microvilli and other membrane protrusions has implicated CD133 in the organization and dynamics of the plasma membrane (Marzesco et al., 2005; Marzesco et al. 2009; Corbeil et al., 2010; Thamm et al., 2019; Pleskač et al., 2024). Importantly, CD133 is not restricted to the plasma membrane and has also been detected in intracellular endosomal compartments, including ILVs within MVBs (Karbanová et al., 2025). These observations place CD133 at the interface between plasma membrane and endosomal membrane organization and raise the possibility that it may contribute to the spatial organization of EV-associated components.

Our previous study identified CD133 as a regulator of EV production and cargo composition in basal-like triple-negative breast cancer (BL-TNBC). We showed that CD133 localizes to plasma membrane protrusions and cholesterol-dependent lipid rafts, where it contributes to the maintenance of membrane architecture and promotes EV release. CD133 was selectively incorporated into tumor-derived EVs and influenced their molecular cargo, including the enrichment of the pro-angiogenic factor CD105/Endoglin. Functionally, CD133-positive EVs promoted endothelial tubulogenesis through a CD105-dependent mechanism. These findings established a link between CD133-dependent membrane organization, EV production and selective cargo incorporation (Gomez-Duro et al., 2026). However, whether CD133-dependent membrane organization is associated with the emergence of distinct EV subpopulations defined by their molecular marker profiles remained unclear.

CD63 is a tetraspanin widely used as an EV-associated marker and is frequently associated with exosomes, reflecting its enrichment in late endosomal compartments, including MVBs, although also being present at the plasma membrane (Pols and Klumperman, 2009; Mathieu et al., 2021). Its distribution between intracellular endosomal compartments and the cell surface makes CD63 a useful protein for investigating the spatial organization of EV-associated components within donor cells. Importantly, however, the presence of CD63 in an EV does not by itself establish its route of biogenesis, as CD63 can be associated with both endosomal and plasma membrane compartments (Pols and Klumperman, 2009; Mathieu et al., 2021). Examining its spatial distribution within the donor cell may therefore provide insight into the relationship between the spatial organization of EV-associated proteins and the heterogeneity of released EV populations.

In our previous study, CD9 was detected in plasma membrane regions associated with CD133-positive domains in BL-TNBC cells (Gomez-Duro et al., 2026). These observations further suggested that CD133-dependent membrane organization may influence the spatial distribution of EV-associated proteins at the plasma membrane. We therefore focused on CD63 as a complementary marker present at both the plasma membrane and in endosomal compartments, enabling us to examine its spatial distribution across these cellular locations.

In the present study, we investigated whether CD133 expression promotes plasma membrane remodeling and whether this remodeling is associated with changes in CD63 distribution and EV heterogeneity. Using complementary TNBC models with different levels of CD133 expression, including loss-, rescue- and gain-of-function conditions, we combined transmission electron microscopy (TEM), quantitative immunogold electron microscopy (IEM), quantitative morphometric analysis of plasma membrane architecture and intracellular MVB compartments, and nano flow cytometry (nFCM) to examine the relationship between CD133, membrane organization, the spatial distribution of EV-associated proteins, and EV heterogeneity.

We hypothesized that CD133 acts not only as a regulator of EV production, but also as a spatial organizer of EV-associated membrane domains. Specifically, we proposed that CD133-dependent remodeling of the plasma membrane contributes to the spatial organization of EV-associated components and is associated with the emergence of heterogeneous CD133/CD63 EV populations.

By integrating analyses of plasma membrane architecture, CD63 distribution, EV heterogeneity and intracellular MVB morphology, this study investigates how CD133-associated membrane remodeling relates to EV marker heterogeneity and intracellular membrane organization in TNBC cells.

## MATERIAL AND METHODS

### Cell Culture

Cell lines were obtained from the American Type Culture Collection (ATCC, Manassas, VA, USA). MDA-MB-231 and MDA-MB-468 cells were cultured in DMEM, high glucose, GlutaMAX™ Supplement (Thermo Fisher Scientific), supplemented with 10% FBS (Gibco), 5% Pen/Strep (Gibco). HCC-70 cells were maintained in RPMI-1640 GlutaMAX™ medium (Sigma-Aldrich) supplemented with 1.5 g/L sodium bicarbonate, 10 mM HEPES, and 1 mM sodium pyruvate (all from Thermo Fisher Scientific), 10% FBS, and 5% Pen/Strep. Genetically modified MDA-MB-231 and MDA-MB-468 cell lines, including MDA-MB-231 CD133-GFP-overexpressing (231-OE), MDA-MB-468 CD133-knockout (468-KO), MDA-MB-468 CD133-knockout CD133-GFP overexpressing (468-rescue), MDA-MB-468 CD133-GFP overexpressing cells (468-OE), were maintained in complete DMEM medium under puromycin selection (2 µg/mL, Thermo Fisher).

All cells were incubated at 37°C in a humidified atmosphere with 5% CO. Upon acquisition, cells were expanded and frozen in aliquots stored in vapor-phase liquid nitrogen. Cell stocks were used up to passage 20. All cultures tested negative for Mycoplasma by PCR.

### Antibodies

The following antibodies were used for WB or IEM: rabbit (rb) anti-Syntenin-1 (Abcam ab133267; 1:1.000 WB); ms monoclonal anti-TSG101 (Genetex GTX70255; 1:1.000 WB); ms anti-human CD81 monoclonal antibody (Santa Cruz sc-166029; 1:50 WB); ms anti-CD9 (Sigma-Aldrich, clone MM2/57 CLB 162; 1:1.000 WB); ms monoclonal CD63 (MBL D263-3, 1:200 WB) or (Thermo Fisher Scientific MA1-35272; 1:50 IEM); ms anti-Alix (Abnova H00010015-B01P; 1:1.000 WB); rb anti-ApoB (Abcam ab312318; 1:1.000 WB); rb anti-hSP70 (Proteintech 10995-1-AP; 1:1.000 WB); rb anti-CD133 clone CD133 D2V8Q (Cell Signaling 64326; 1:1.000 WB); anti-CD133/1 (Miltenyi Biotec 130-113-670; 1:1.000 WB, 1:50 IEM); rb anti-GAPDH (Sigma G9545; 1:10.000 WB). Secondary antibodies coupled to HRP-conjugated goat polyclonal antibodies to rabbit IgG (Abcam ab6721) or to mouse IgG (Abcam ab6789) were used at 1:5.000. Protein A-gold particles (10 nm; Cell Microscopy Center, Department of Cell Biology, Utrecht University) were used for immunogold labelling.

### Plasmids

The plasmids used for CRISPR/Cas9-mediated CD133 (PROM1) knockout in MDA-MB-468 cells, including the dual-guide lentiviral plasmid pLV[2CRISPR]-hCas9 (VectorBuilder, VB221117-1157xvx) and the corresponding scramble control, were previously described in Gomez-Duro et al. (2026). Plasmids were prepared as previously described.

### Generation of CD133 KO Cells by CRISPR/Cas9

CD133 knockout (CD133 KO) MDA-MB-468 cells were generated using the lentiviral CRISPR/Cas9 system described previously Gomez-Duro et al. (2026). Briefly, MDA-MB-468 cells were transduced with lentiviral particles carrying the dual-guide CRISPR/Cas9 vector targeting the human *PROM1* gene and selected with puromycin, using the conditions previously described. A scramble control cell line was generated in parallel using the corresponding non-targeting control vector.

### Lentiviral CD133-GFP Expression and Rescue

CD133-GFP expression was introduced into CD133-KO MDA-MB-468 (468-KO) cells to generate rescue cells and into parental MDA-MB-468 (468-OE) and MDA-MB-231 (231-OE) cells to generate CD133-overexpressing cells. The full-length human *PROM1* coding sequence fused to GFP, under the control of the CMV promoter, was derived from the pCD133-GFP vector kindly provided by D. Corbeil (Reichert et al., 2016) and cloned into a lentiviral vector, as previously described (Gomez-Duro et al., 2026). Lentiviral particles were produced by transient transfection of HEK293T cells with the CD133-GFP construct and the packaging plasmids psPAX2 and pMD2.G, using TransIT-293 Transfection Reagent (Mirus Bio) according to the manufacturer’s instructions. Viral supernatants were collected 48 h after transfection, centrifuged at 350 × *g* for 10 min, filtered through a 0.45-µm filter and applied to the target cells. GFP-positive cells were isolated by fluorescence-activated cell sorting (FACS), expanded under standard culture conditions, and CD133-GFP expression was confirmed by Western blot.

### Serum EVs-depleted Medium

To eliminate EVs from FBS, 250 mL of RPMI or DMEM containing 20% FBS was ultracentrifuged at 100,000 × g for 18 hours at 4°C using a Ti45 rotor (Beckman Coulter). After ultracentrifugation, the EV-depleted supernatant was collected and supplemented with 5% Pen/Strep and an additional 250 mL of fresh RPMI or DMEM to obtain the final working concentration. The mixture was then filtered through a 0.22 µm Stericup sterile vacuum filtration system (Millipore Sigma). This FBS EVs-depleted medium was used for all EV-related experiments.

### EVs Isolation by SEC

Cells were cultured in EV depleted medium for 24 h. Cell number and viability were assessed at the time of conditioned medium collection, and cell viability was consistently above 90% across all experimental conditions. Conditioned medium was harvested by pelleting cells at 300 × g centrifugation, at 4°C. The supernatant was further centrifuged at 2,000 × g for 20 minutes at 4°C to remove cell debris. The resulting supernatant was concentrated using Centricon Plus-70 centrifugal filters (Millipore, 10 kDa molecular weight cutoff) at 2,000 × g at 4°C until the final volume was reduced to less than 500 µL. The concentrated CM was then applied to qEV original SEC columns (Izon, SP5), which allowed optimal recovery of vesicles ranging from 35 to 350 nm. The void volume (fractions 1-6, totaling 3 mL) was discarded, and fractions 7-11 (2.5 mL), enriched in EV, were collected. These fractions were then concentrated using Amicon Ultra-15 centrifugal filters (Millipore, 10 kDa cutoff) at 3,320 × g at 4°C. Aliquots of the final EV preparations were stored at -80°C until use.

### Single-particle Characterization of EVs

#### Nanoparticle Tracking Analysis (NTA)

NTA was performed using the ZetaView PMX-120 system (Particle Metrix, software v8.04.02) at 22°C. Instrument settings were sensitivity 75, shutter speed 75, and frame rate 30 fps. Two independent dilutions of each sample were analyzed across 11 positions, with two cycles per position.

#### Nano-Flow Cytometry (nFCM)

For high-resolution single-particle analysis, nFCM was carried out using the NanoAnalyzer (NanoFCM, S/N FNAN30E22101948). The instrument used single-photon-counting modules to detect side scatter (SSC, 488/10 nm), green fluorescence (525/40 nm), and far-red fluorescence (670/30 nm). Calibration for particle concentration was done using 250 nm fluorescent silica QC beads, and particle sizing was calibrated with a standard mix of silica beads (68, 91, 113, and 155 nm; S16M-Exo beads). Samples were diluted in 0.2 µm-filtered 1× PBS to reach a final concentration of 2,000–12,000 particles/min, measured at 1 kPa pressure. Data were recorded using NanoFCM Professional 2.0 software and exported as.fcs files for analysis in FlowJo v10.8.1.

### EV Surface Immunolabeling and Nano-Flow Cytometry Analysis

EV surface marker analysis by nano-flow cytometry (nFCM) was performed following the protocol described by Nevo et al. 2026 (Nevo et al., 2026). SEC-purified EVs were diluted in PBS to 2-3 × 10¹ particles/mL and incubated with fluorophore-conjugated antibodies for 30 min at 4°C. Samples were subsequently analyzed directly or subjected to SEC-based purification to remove unbound antibodies before acquisition. Antibody-only controls were included to assess background fluorescence and establish staining specificity. Gates were defined using unstained EV and PBS preparations. Data were acquired on a NanoAnalyzer (NanoFCM) and analyzed using NanoFCM Professional software and FlowJo v10.8.1.

### Multiplex Bead-Based Flow Cytometry for EV Surface Marker Profiling

EVs isolated from TNBC cell lines via SEC were characterized using bead-based multiplex FC (MACSPlex Exosome Kit, human; Miltenyi Biotec). EV input was standardized based on particle concentration determined by NTA. A total of 1 × 10 EVs were diluted in MACSPlex buffer to a final volume of 120 µL and incubated overnight at RT with 10 µL of MACSPlex Exosome Capture Beads on an orbital shaker, protected from light. Following incubation, samples were washed and stained for 1 hour at RT with APC-conjugated detection antibodies against CD9, CD63, and CD81 (provided in the kit), used as a combined antibody cocktail or, where indicated, individually for CD63 or CD9 detection. Flow cytometric analysis was performed on a MACSQuant Analyzer 10 (Miltenyi Biotec) following the manufacturer’s recommendations. Data were analyzed using FlowJo software (v10, FlowJo LLC). Median fluorescence intensity (MFI) values were normalized against the corresponding blank control included in each experiment to correct for background fluorescence. Among the 37 markers included in the assay, only selected markers relevant to the study objectives are reported.

### Western Blot Analysis (WB)

Cells were harvested and washed twice with cold PBS. Approximately 1 × 10 cells were pelleted by centrifugation at 300 × g for 5 minutes at 4°C and resuspended in 200 µL of lysis buffer containing 50 mM Tris-HCl (pH 7.5), 150 mM NaCl, 10% (v/v) glycerol, 5 mM EDTA, and 1% (v/v) Triton X-100, supplemented with a protease inhibitor cocktail (Sigma-Aldrich; P8340). Lysates were incubated on ice for 20 minutes with gentle agitation, followed by centrifugation at 16,000 × g for 20 minutes at 4°C to remove insoluble material. The resulting clarified supernatants were quantified using the Pierce™ BCA Protein Assay Kit (Thermo Fisher Scientific; catalog no. 23225). Both whole-cell lysates and EV were mixed with Laemmli sample buffer (BioRad; catalog no. 1610747), with or without 400 mM DTT (BioRad; catalog no. 1610747), and denatured by boiling for 5 minutes at 95°C or for 10 minutes at 70°C, respectively. Proteins were separated on NuPAGE (4-12%) Bis-Tris gels (Invitrogen) and transferred onto nitrocellulose membranes (Millipore). Membranes were blocked for 1 hour at RT in PBS/ Tween-20 0.1% (PBS/T) with 5% non-fat dry milk, incubated overnight at 4°C with the indicated primary antibodies diluted in the same blocking solution. After three washes with PBS/T, membranes were incubated with HRP-conjugated secondary antibodies, followed by additional washes in PBS/T. Immunoreactive bands were visualized using Clarity™ Western ECL SubstrateBio-Radad, catalog no. 170-5060) and detected using the ChemiDoc Touch Imaging System (Bio-Rad), ensuring exposure within the linear detection range. Band intensities were quantified by densitometry using the “Analyze Gels” tool in ImageJ (https://imagej.net/ij/).

### Transmission Electron Microscopy (TEM)

Cellular ultrastructure and immunogold labeling of cells were performed using established protocols adapted from Hurbain et al. 2017.

#### Cellular ultrastructure

TNBC cell monolayers were fixed in a solution of 2.5% glutaraldehyde (Electron Microscopy Sciences; catalog no. 16220) in 0.1 M cacodylate buffer pH 7.2 (EMS, Cat #12300) for 24 hours at 4°C. Post-fixation was performed using 1% osmium tetroxide (EMS, Cat #19150), supplemented with 1.5% potassium ferrocyanide,(Sigma #702587) in 0.1 M cacodylate buffer pH 7.2 45 min at 4°C, followed by graded ethanol dehydration and embedding in resin (EMbed 812 Embedding Kit, EMS cat #14120). Using an ultramicrotome UC6 (Leica)with a diamond knife (DiATOME), cells sections were collected on copper grids (Delta microscopy), mounted and contrasted with uranyl acetate (UA) and lead citrate (Delta Microscopies, ref 11300). Imaging was performed using a Tecnai Spirit G2 transmission electron microscope (ThermoFisher Scientific) operated at 80 kV, equipped with a 4k CCD camera (Gatan OnView 4k x 4k).

#### Immunogold labeling of cryosections

TNBC cells at over 80% confluence were fixed overnight at 4°C in 0.1 M PB containing 0.25% glutaraldehyde (Electron Microscopy Sciences; catalog no. 16220) and 2% PFA (Electron Microscopy Sciences; catalog no. 15710). After fixation, cells were washed three times with PBS and incubated for 10 minutes in PBS with 0.15% glycine to neutralize free aldehydes. Cells were scraped, centrifuged, and resuspended in 12% gelatin, followed by incubation at 37°C and a last centrifugation. Embedded pellets were solidified on ice, cut in 1mm³ cubes, and cryoprotected overnight at 4°C in 2.3 M sucrose. Blocks were mounted on steel pins, and rapidly frozen in liquid nitrogen. Ultrathin sections (∼90 nm) were prepared on a cryo-ultramicrotome (Leica UC7) at −110°C, collected on formvar- and carbon-coated grids, and stored at 4°C. Immunolabeling was performed by incubating grids in 2% gelatin, washing with PBS-glycine, blocking with 1% BSA, and applying primary antibodies diluted in PBS/1% BSA for 1 hour at RT. After washing, mouse primary antibodies were bridged with an anti-mouse IgG before incubation with protein A-gold. Following final washes, grids were fixed in 1% glutaraldehyde (Electron Microscopy Sciences; catalog no. 16220), rinsed, and contrast-stained with 2% UA and methylcellulose at pH 4.

Cell samples were examined using a Tecnai Spirit G2 transmission electron microscope (ThermoFisher Scientific) operated at 80 kV and equipped with a 4k CCD camera (On View 4k x 4k GATAN).

### Image analysis

#### Cell and membrane protrusion analysis

A Cellpose model (Stringer et al., 2021) was trained to segment membrane protrusions, starting from the cyto2 model. Twenty images were manually annotated for training, using SAMJ (Garcia-Lopez-de-Haro et al., 2026) for initial segmentation, followed by manual correction where required. Model training was performed using default parameters.

For each cell, two regions of interest (ROIs) were generated by manual segmentation: one excluding membrane protrusions and one including the cell body and membrane protrusions. These ROIs were used to quantify cell and membrane protrusion features, including protrusion number, protrusion density, protrusion length, and the proportion of the analyzed membrane occupied by protrusions.

Protrusion length was estimated using a custom Fiji macro. Briefly, random coordinate points were sampled along the cell contour, and points corresponding to the protrusion-containing ROI were retained. For each retained point, the normal to the cell contour was calculated from the coordinates of the two adjacent contour points. A line was then generated from the reference point along the outward normal until it intersected the boundary of the ROI including the cell body and membrane protrusions. The distance between the reference point and the intersection was used as an estimate of protrusion length. Fifteen measurements were performed per cell and averaged to obtain the mean protrusion length.

#### MVB analysis

MVBs were segmented using the SAMJ plugin available in Fiji (Garcia Lopez-de-Haro et al., 2026). Based on these segmentations, a custom Fiji macro was used to extract MVB morphological features, including area, circularity, aspect ratio (AR), and size distribution. The macro also required manual segmentation of the cell contour to determine MVB density (number of MVBs/µm²), the percentage of cell area occupied by MVBs, and the MVB burden index.

### Statistical Analysis

All statistical analyses were conducted using GraphPad Prism version 10.5.0. Quantitative data are presented as mean ± standard error of the mean (SEM). For comparisons between two groups, the Student’s t-test was used. For comparisons involving more than two groups, either one-way ANOVA or the Kruskal-Wallis test was applied, depending on data distribution. Post hoc analyses were performed using Tukey’s or Dunn’s multiple comparisons tests, as appropriate. For experiments involving two independent variables, two-way ANOVA followed by Tukey’s multiple comparisons test was used. A p-value < 0.05 was considered statistically significant. Statistical significance was defined as follows: p < 0.05 (*), p < 0.01 (**), p < 0.001 (***), and p < 0.0001 (****).

## RESULTS

### CD133-expressing TNBC cells display a highly protrusive plasma membrane

To determine whether CD133 expression is associated with differences in plasma membrane architecture, we first examined the ultrastructure of three TNBC cell lines with distinct CD133 expression levels, as previously characterized (Gomez-Duro et al., 2026) : CD133-low MDA-MB-231 cells representative of the mesenchymal-like (MES) subtype, and CD133-high MDA-MB-468 and HCC-70 cells, both representative of the basal-like (BL) subtype. TEM revealed marked differences in plasma membrane morphology between the three cell lines (Figure 1A). MDA-MB-231 cells generally displayed a relatively flat plasma membrane, with occasional protrusive structures. In contrast, both MDA-MB-468 and HCC-70 cells exhibited numerous prominent membrane protrusions extending from the cell surface.

**Figure 1.**
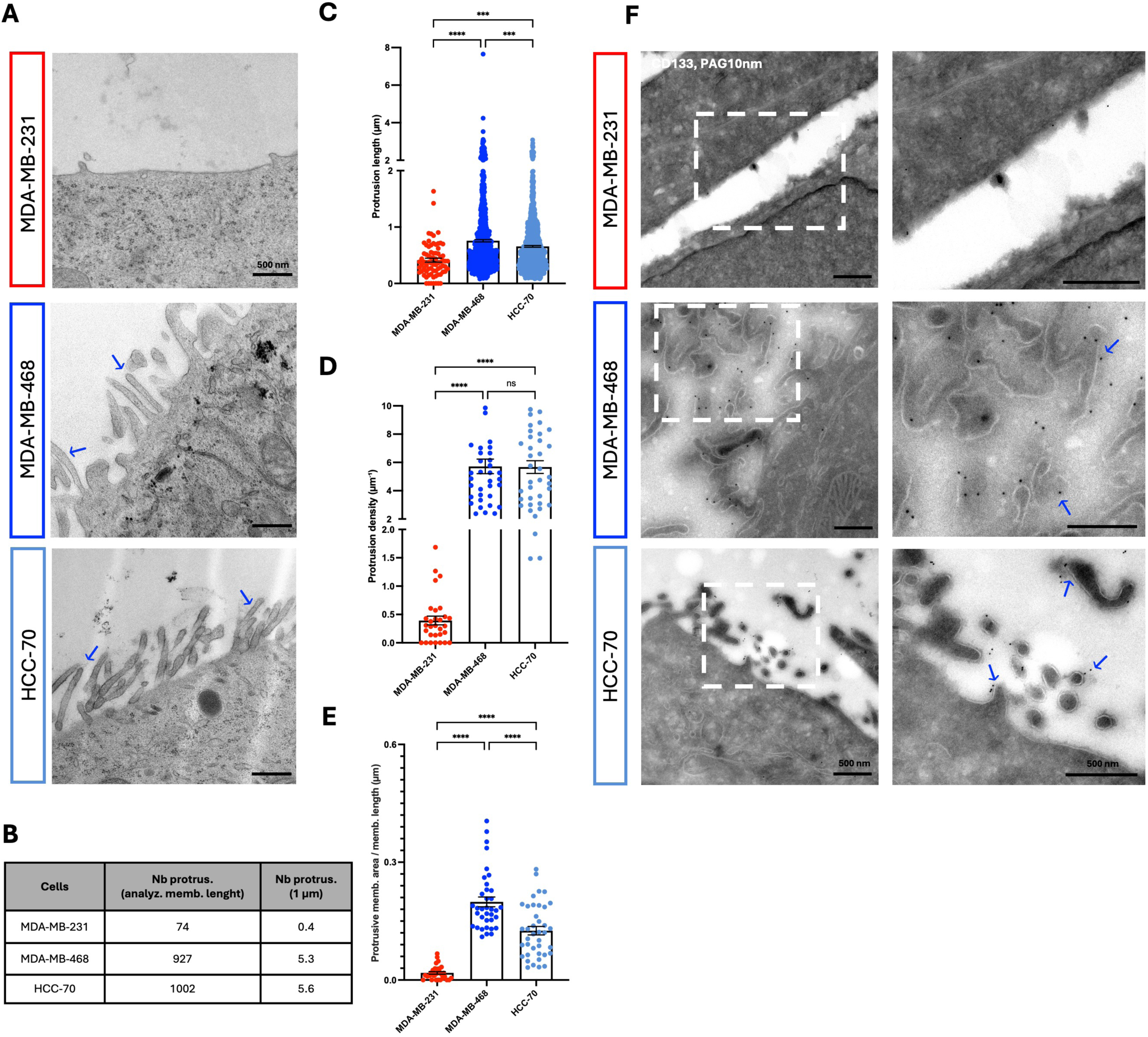
CD133-expressing TNBC cells display a highly protrusive plasma membrane. **A**. Representative TEM micrographs of TNBC cells (MDA-MB-231, MDA-MB-468 and HCC-70). Blue arrows indicate membrane protrusions. Scale bar: 500 nm. **B.** Quantification of the total number of protrusions couted along the analyzed membrane and the corresponding protrusion density (protrusions/µm of membrane) in the three TNBC cell lines. **C**. Distribution of protrusion length (µm) for each TNBC cell line. Data are presented as mean ± SEM from 3 independent experiments. Statistical significance was assessed by one-way ANOVA. \*\*\*\**p* < 0.0001; \*\*\**p* = 0.0006 (MDA-MB-231 *vs*. HCC-70) and \*\*\**p* = 0.0002 (MDA-MB-468 *vs*. HCC-70). **D**. Individual values of protrusion density (protrusions/µm of membrane) for each TNBC cell line, with individual values shown. Data are presented as mean ± SEM from 3 independent experiments. Statistical significance was assessed by one-way ANOVA. \*\*\*\**P* < 0.0001 and ns = 0.9962 (MDA-MB-468 *vs.* HCC70). **E.** Ratio of protrusive membrane area relative to membrane length (µm) for each TNBC cell line. Data are presented as mean ± SEM from 3 independent experiments. Statistical significance was assessed by one-way ANOVA. \*\*\*\**p* < 0.0001. **F.** Representative ultrathin cryosections of each TNBC cell line immunolabeled with anti-CD133-PAG (10 nm). Blue arrows indicate CD133 positive staining localized within membrane protrusions. Right: magnification of the white boxed regions. Scale bars are indicated in each image.

To quantify these differences, we developed a dedicated image-analysis macro to systematically measure protrusion number, length and area from TEM images (Figure 1B-E). Protrusion length was significantly higher in both CD133-expressing MDA-MB-468 and HCC-70 cells than in MDA-MB-231 cells (Figure 1C). Protrusion density was markedly higher in MDA-MB-

468 and HCC-70 cells than in MDA-MB-231 cells, reaching approximately 5.3 and 5.6 protrusions/µm of membrane, respectively, compared with 0.4 protrusions/µm in MDA-MB-231 cells (Figure 1B, D). Accordingly, the proportion of plasma membrane occupied by protrusive structures was markedly greater in MDA-MB-468 and HCC70 cells than in MDA-MB-231 cells, with values of approximately 20%, 13%, and 2%, of the analyzed membrane respectively (Figure 1E).

To determine whether CD133 itself was localized within these protrusive membrane domains, we performed IEM in the three cell lines. CD133 labeling was not detected at the plasma membrane of MDA-MB-231 cells, consistent with our previous characterization of this cell line (Gomez-Duro et al., 2026). In contrast, CD133 labeling was readily detected in both MDA-MB-468 and HCC-70 cells and was observed within membrane protrusions (Figure 1F). The localization of CD133 to protrusions in MDA-MB-468 cells is consistent with our previous study (Gomez-Duro et al. 2026), while the present analysis extends this observation to HCC-70 cells.

Together, these ultrastructural analyses establish a strong association between CD133 expression and a protrusive plasma membrane architecture in TNBC cells.

### CD133 expression promotes plasma membrane protrusion

To directly assess the contribution of CD133, we generated stable CD133-GFP-expressing cells in the previously characterized CD133-KO MDA-MB-468 background (Gomez-Duro et al., 2026, 468-rescue cells) and in MDA-MB-231 cells (231-OE cells), resulting in rescue and OE models, respectively. As shown in Figure S1A-E, 468-KO cells displayed reduced protrusion length, density, and protrusive membrane area compared with parental and scramble control cells, whereas parental and scramble control cells showed comparable protrusion morphology. Immunogold labeling further confirmed the presence of CD133 within membrane protrusions in scramble control cells, but not in 468-KO cells (Figure S1F). CD133-GFP was also expressed in MDA-MB-468 cells retaining endogenous CD133 expression to assess the effect of increased CD133 levels (468-OE cells). WB analysis confirmed the expected CD133 expression profile across the different experimental conditions (Figure 2A). Endogenous CD133 was detected in MDA-MB-468 control cells but not in CD133-KO cells (468-KO) or MDA-MB-231 cells, whereas CD133-GFP was detected in the rescue (468-rescue), OE (231-OE) and MDA-MB-468-OE (468-OE) conditions. Quantification of CD133 relative to GAPDH confirmed the expected differences in CD133 abundance between the different conditions (Figure 2B).

**Figure 2.**
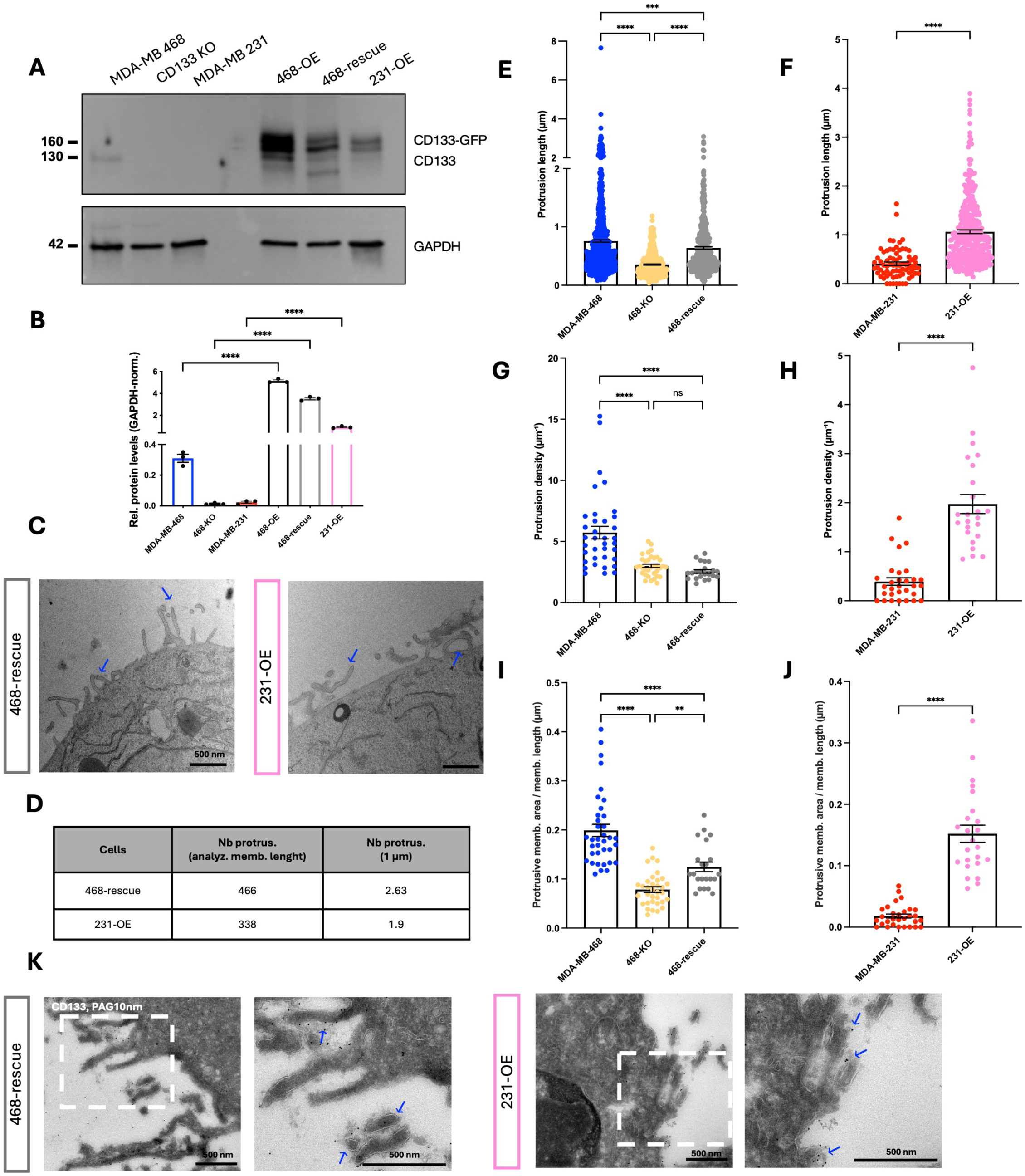
CD133 expression is sufficient to promote plasma membrane protrusion. **A.** Western blot analysis of cell lysates (10 µg) from parental MDA-MB-468, 468-KO and MDA-MB-231 and 468-OE, 468-rescue and 231-OE cells probed for CD133 and GAPDH as a loading control. **B.** Quantification of CD133 protein levels in cell lysates normalized to GAPDH. Data are presented as mean ± SEM from 3 independent experiments, analyzed by one-way ANOVA. \*\*\*\**P* < 0.0001. **C.** Representative TEM micrographs of rescue and 231-OE cells. Blue arrows indicate membrane protrusions. Scale bar: 500 nm. **D.** Quantification of the total number of protrusions couted along the analyzed membrane and the corresponding protrusion density (protrusions/µm of membrane) in 468-rescue and 231-OE cells. **E-F**. Distribution of protrusion length (µm) in (**E**) MDA-MB-468, 468-KO, and 468-rescue cells., and (**F**) MDA-MB-231 and 231-OE cells. Statistical significance was assessed by one-way ANOVA for MDA-MB-468, CD133 KO, and rescue cells and by an unpaired t-test for MDA-MB-231 and 231-OE cells. \*\*\*\**p* < 0.0001 (MDA-MB-468 *vs*. 468-KO, 468-KO *vs.* 468-rescue, MDA-MB-231 *vs.* 231-OE); \*\*\**p* = 0.0001 (MDA-MB-468 *vs*. rescue). **G-H.** Individual values of protrusion density (protrusions/µm of membrane) in (**G**) MDA-MB-468, CD133-KO, and rescue cells, and (**H**) in MDA-MB-231 and 231-OE cells. Statistical significance was assessed by one-way ANOVA for MDA-MB-468, CD133 KO, and rescue cells and by an unpaired t-test for MDA-MB-231 and 231-OE cells. \*\*\*\**p* < 0.0001 (MDA-MB-468 *vs.* 468-KO; MDA-MB-468 *vs.* 468-rescue and MDA-MB-231 *vs.* 231-OE); ns = 0.6915 (CD133 KO *vs.* rescue). **I-J.** Ratio of protrusive membrane area relative to membrane length (µm) in (**I**) MDA-MB-468 parental, CD133-KO, and rescue cells, and (**J**) in MDA-MB-231 and 231-OE cells. Statistical significance was assessed by one-way ANOVA for MDA-MB-468, CD133 KO, and rescue cells and by an unpaired t-test for MDA-MB-231 and 231-OE cells. \*\*\*\**p* < 0.0001 (MDA-MB-468 *vs.* 468-KO; MDA-MB-468 *vs.* 468-rescue and MDA-MB-231 *vs.* 231-OE); \*\**p* = 0.0093 (468-KO *vs.* 468-rescue). **K.** Representative ultrathin cryosections of 468-rescue and 231-OE cells immunolabeled with anti-CD133-PAG (10 nm). Blue arrows indicate CD133 staining localized in membrane protrusions. Right: magnification of the white boxed region. Scale bar: 500 nm.

We next examined plasma membrane ultrastructure by TEM in the 468-rescue and 231-OE models (Figure 2C). Restoration of CD133 expression in 468-KO cells was associated with the reappearance of prominent membrane protrusions (compare Figure S1 to Figure 2C), whereas expression of CD133-GFP in MDA-MB-231 cells, which normally display a relatively flat plasma membrane (Figure 1A), resulted in the appearance of numerous membrane protrusions (Figure 2C).

We then quantified protrusion length, protrusion density, and protrusive membrane area separately in the two experimental models. In the MDA-MB-468 series, comparison of parental, 468-KO, and 468-rescue cells showed that CD133 loss was associated with reduced protrusion length, protrusion density, and protrusive membrane area, whereas CD133 restoration partially recovered these morphological features (Figure 2D, E, G, I). Protrusion density was 5.3 protrusions/µm in parental MDA-MB-468 cells and 2.63 protrusions/µm in rescue cells (Figures 1B, 2D). These findings are consistent with our previous characterization of 468-KO cells (Gomez-Duro et al., 2026). In the MDA-MB-231 model, CD133 expression (231-OE) markedly increased protrusion length, protrusion density, and protrusive membrane area compared with parental cells (Figure 2F, H, J), with protrusion density increasing from 0.4 to 1.9 protrusions/µm (Figure 2D), corresponding to an approximately 5-fold increase.

Finally, IEM confirmed the localization of CD133 within the protrusive membrane domains in both experimental models (Figure 2K). CD133 immunogold labeling was detected within membrane protrusions in 468-rescue cells and 231-OE cells.

Together, these loss-, rescue-, and gain-of-function experiments support a role for CD133 in promoting protrusive plasma membrane organization. In particular, CD133 expression in MDA-MB-231 cells, which lack detectable endogenous CD133, induced a pronounced protrusive phenotype.

### CD133 expression redistributes CD63 toward protrusive plasma

We next investigated whether the protrusive membrane domains associated with CD133 expression were characterized by a distinct distribution of the EV-associated tetraspanin CD63. Because CD63 is present both at the plasma membrane and in intracellular endosomal compartments, including MVBs (Pols and Klumperman, 2009; Mathieu et al., 2021), we first examined its spatial distribution at the plasma membrane of MDA-MB-231 control and 231-OE cells by quantitative IEM (Figure 3A). Additional representative IEM images and cell-level raw measurements are provided in Figure S2.

**Figure 3.**
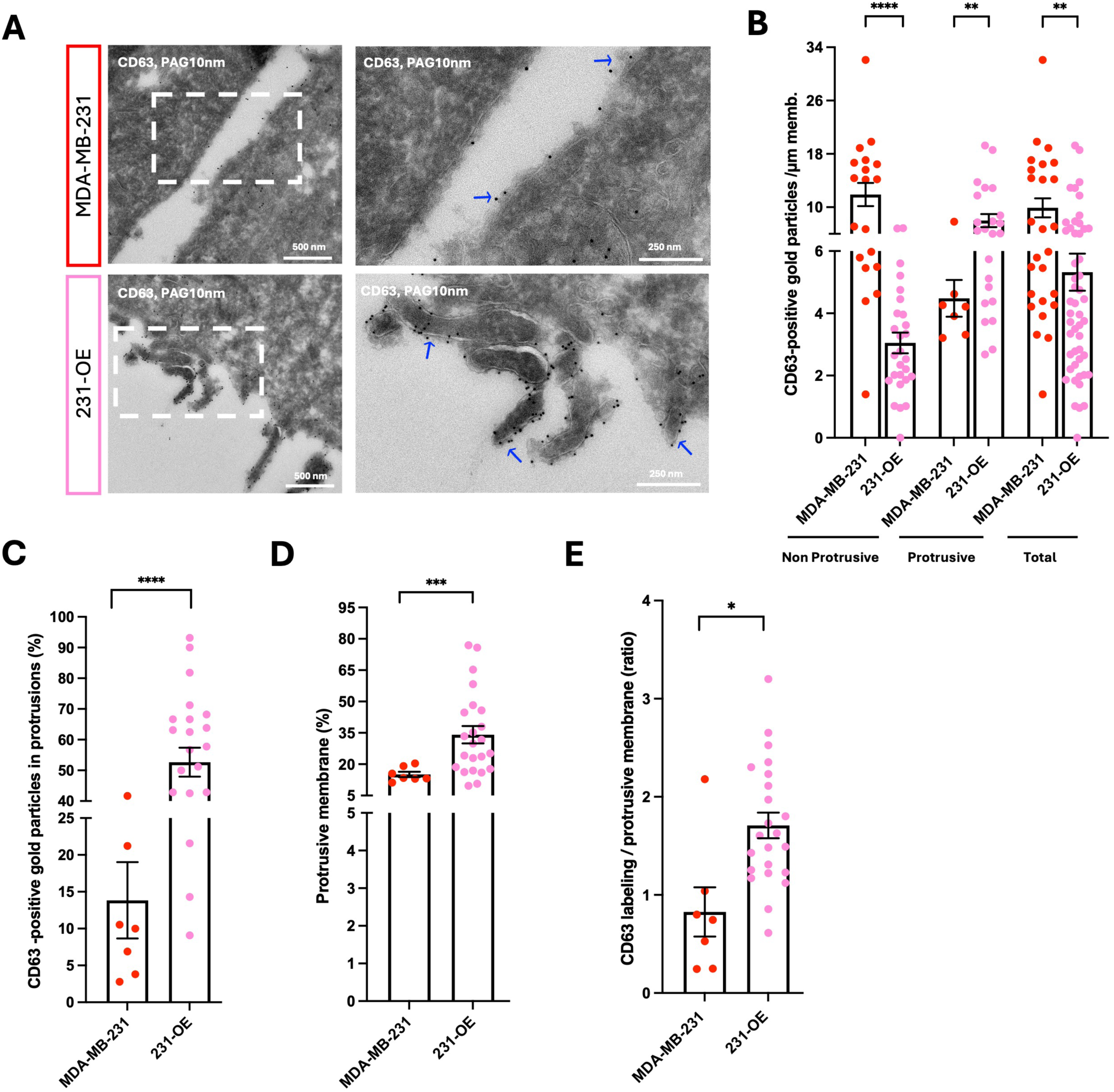
CD133 expression redistributes CD63 toward plasma membrane protrusions in MDA-MB-231 cells. **A**. Representative immunogold electron microscopy (IEM) images showing CD63 localization at the plasma membrane of MDA-MB-231 and 231-OE cells. Blue arrowheads indicate CD63-positive gold particles associated with protrusive membrane domains. Right: magnification of the white boxed region. Scale bars are indicated in each image. **B**. Quantification of CD63 immunogold particle density in non-protrusive and protrusive plasma membrane domains in 231 and 231-OE cells. Data are presented as mean ± SEM, analyzed by unpaired t-test for MDA-MB-231 and 231-OE cells. \*\*\*\**p* < 0.0001 (MDA-MB-231 Non-Protrusive *vs*. 231-OE Non-Protrusive). \*\**p* = 0.0048 (MDA-MB-231 Protrusive *vs*. 231-OE Protrusive) and \*\**p* = 0.0058 (MDA-MB-231 Total *vs*. 231-OE Total). **C**. Percentage of CD63 immunogold particles localized within protrusive membrane domains in 231 and 231-OE cells. Data are presented as mean ± SEM, analyzed by unpaired t-test for MDA-MB-231 and 231-OE cells. \*\*\*\**p* < 0.0001. **D.** Percentage of the total analyzed plasma membrane length occupied by protrusions in 231 and 231-OE cells. Data are presented as mean ± SEM, analyzed by unpaired t-test for MDA-MB-231 and 231-OE cells. \*\*\**p* = 0.0002. For all quantifications, each dot represents one cell; 231 n = 19 cells. 231-OE n = 27 cells. **E.** Relative representation of CD63 labeling within membrane protrusions, calculated as the ratio between the percentage of CD63-positive gold particles localized to protrusions and the percentage of plasma membrane occupied by protrusions, in MDA-MB-231 and 231-OE cells. Data are presented as mean ± SEM, analyzed by unpaired t-test for MDA-MB-231 and 231-OE cells. \**p* = 0.0116. 231 n = 19 cells. 231-OE n = 27 cells.

In MDA-MB-231 cells, CD63 labeling was detected predominantly along non-protrusive regions of the plasma membrane. The average CD63 density was 11.9 gold particles/µm on non-protrusive membrane, compared with 4.5 gold particles/µm within protrusive membrane domains (Figure 3B). In 231-OE cells, CD63 density on non-protrusive membrane decreased to 3 gold particles/µm, whereas CD63 labeling density was igher within protrusions than along non-protrusive membrane, with an average density of 8 gold particles/µm (Figure 3B). Considering the plasma membrane as a whole, the average CD63 density was 9.9 gold particles/µm in MDA-MB-231 cells and 5.3 gold particles/µm in 231-OE cells.

We then quantified the fraction of membrane-associated CD63 labeling located within protrusions. In MDA-MB-231 cells, 13.8 % of CD63-positive gold particles were associated with protrusive membrane, close to the 15.1% of the analyzed plasma membrane occupied by protrusions (Figure 3C, D). Following CD133 expression, this relationship was markedly altered: 52.7% of CD63-positive gold particles were localized to protrusions, compared with 34.1 % of the analyzed plasma membrane occupied by protrusions (Figure 3C, D). To account for differences in protrusion coverage, we calculated the ratio between the fraction of CD63 labeling in protrusions and the fraction of membrane occupied by protrusions. This was 0.83 in control cells and 1.55 in 231-OE cells, corresponding to an approximately 1.9-fold increase in the relative representation of CD63 labeling in protrusions following CD133 expression (Figure 3E). Thus, CD63 labeling approximately reflected the proportion of protrusive membrane in MDA-MB-231 cells, whereas CD133 expression was associated with a disproportionate representation of CD63 labeling within protrusive membrane domains.

### CD133 expression increases particle output and alters EV-associated marker profiles

We next investigated whether CD133-dependent membrane remodeling was accompanied by changes in the EV population released by MDA-MB-231 cells. EVs from control and 231-OE cells were characterized using complementary approaches, including nano-flow cytometry (nFCM), nanoparticle tracking analysis (NTA), WB and multiplex analysis of EV surface markers (MACSPLex EV) (Figure 4).

**Figure 4.**
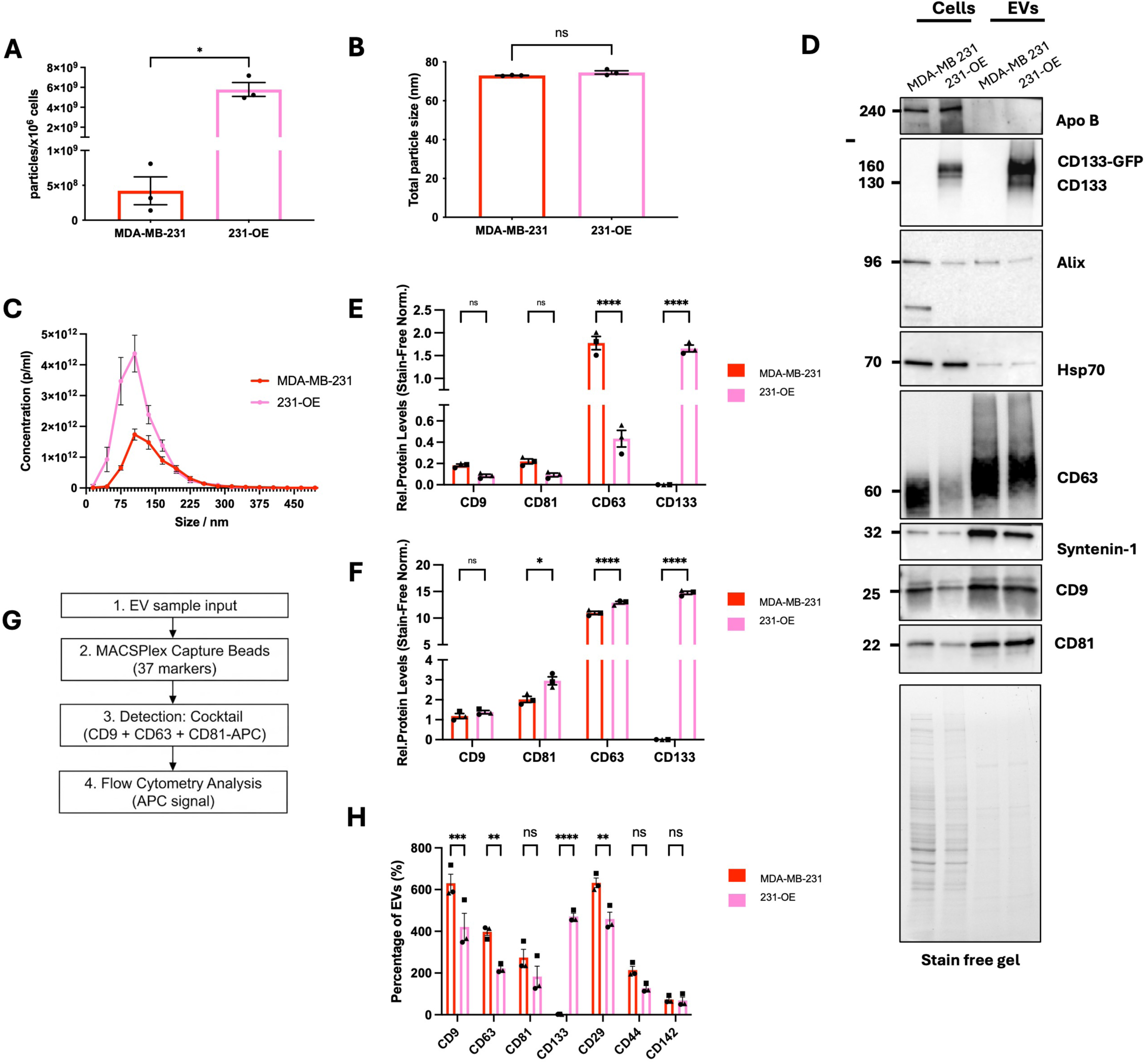
CD133 expression increases EV production and alters EV molecular composition. **A.** Histogram showing EV particles concentration (particles per million cells) in MDA-MB-231 and 231-OE cells, measured by nFCM. Data are presented as mean ± SEM from 3 independent experiments, analyzed by unpaired t-test. \**p* = 0.0115. **B.** Quantification of EV particle size (median value) measured by nFCM in MDA-MB-231 and 231-OE cells. Data are shown as mean ± SEM from 3 independent experiments, analyzed by unpaired t-test. ns = 0.2120. **C.** NTA showing EV from MDA-MB-231 and 231-OE cells (n = 3 independent experiments). The graph displays the concentration of particles (particles/mL) as a function of particle size (nm). Peaks indicate the predominant particle sizes in the MB-231 and 231-OE cells probed for CD133 and canonical EV markers. **E.** Stain free gel image and quantification of protein levels in cell lysates normalized to total protein. Data are presented as mean ± SEM from 3 independent experiments, analyzed by two-way ANOVA. ns = 0.7647 and 0.5614 (CD9 and CD81, respectively); **** *p* < 0.0001 (CD63 and CD133, respectively). **F.** Stain free gel image and quantification protein levels in isolated EVs normalized to total protein. Data are presented as mean ± SEM from 3 independent experiments, analyzed by two-way ANOVA. ns = 0.9403 (CD9); * *p* = 0.0212 (CD81); **** *p* < 0.0001 (CD63 and CD133, respectively). **G.** Schematic illustrating the MACSPlex workflow. EVs were captured by antibody-coated bead populations and detected using APC-labelled antibody cocktail against CD9, CD63 and CD81. **H**. MACSPlex analysis of individual CD9, CD63 and CD8, CD133 and other surface proteins signals detected on EVs, normalized to the blank. Data are presented as mean ± SEM from 3 independent experiments and were analyzed using two-way ANOVA. \*\*\**p* = 0.0004 (CD9); \*\**p* = 0.0027 and 0.0033 (CD63 and CD29, respectively); ns = 0.2827, 0.3369 and > 0.9999 (CD81, CD44 and CD142, respectively); **** *p* < 0.0001 (CD133).

nFCM analysis revealed a significant increase in particle output by 231-OE cells compared with control MDA-MB-231 cells (Figure 4A). Particle output increased from approximately 4.1 × 10^8^ to 6 × 10^9^ particles per 10^6^ cells following CD133 expression, corresponding to an approximately 15-fold increase (Figure 4A). The median particle size did not differ significantly between conditions and was approximately 70 nm (Figure 4B). NTA measurements also showed an increase in particle concentration in the 231-OE condition, from 1.8 × 10^12^ to 4.3 × 10^12^ particles/mL, while the size distribution peaked at approximately 105 nm, in both conditions (Figure 4C). Together, these complementary measurements indicate that CD133 expression in MDA-MB-231 cells was associated with a substantial increase in particle output without a detectable change in the overall size profile.

We next examined the molecular composition of the EV preparations and cell lysates by western blotting (WB) (Figure 4D-F). EV fractions from both conditions displayed EV-associated proteins, including Alix, CD63, Syntenin-1, CD81 and CD9. HSP70 was also detected in both cellular lysates and EV fractions, whereas ApoB was assessed as a marker of potential lipoprotein contamination. Using an anti-CD133 antibody, CD133-GFP was detected in both cell lysates and EVs from 231-OE cells but was not detectable in the corresponding control samples, consistent with the previously reported absence of detectable endogenous CD133 in control MDA-MB-231 cells (Gomez-Duro et al., 2026). Total protein staining by stain-free imaging was used to normalize WB band intensities (Figure 4D-F).

Across the three independent experiments, CD63 signals differed significantly between control and 231-OE samples in both cell lysates and EV preparations, whereas differences in CD9 and CD81 signals did not reach statistical significance in either sample type (Figure 4 E, F). These measurements describe relative protein signals within each sample type and do not establish absolute changes in the cellular abundance or EV loading of individual tetraspanins.

We further characterized EV-associated surface-marker profiles using MACSPlex analysis (Figure 4G, H). EVs were captured using beads coupled to antibodies against specific surface markers, including CD9, CD63, CD81 and CD133, as well as CD29, CD44 and CD142. Captured EVs were then detected using a cocktail of APC-conjugated antibodies against CD9, CD63 and CD81.Background-corrected APC fluorescence signals were expressed as MFI values relative to the corresponding MACSPlex blank control.

Overall, APC signals decreased in the 231-OE condition across several capture-bead populations (Figure 4H). Significant reductions were observed for CD9-capture beads (approximately 35%), CD63-capture beads (approximately 50%) and CD29-capture beads (approximately 30%), whereas changes in the other capture-bead populations did not reach statistical significance. In contrast, a strong APC signal was detected on CD133-capture beads in the 231-OE condition but not in control samples (Figure 4H), consistent with the absence of detectable endogenous CD133 in MDA-MB-231 cells. Because CD9, CD63 and CD81 were detected simultaneously, the APC signals reflect their combined detection on captured EV populations and cannot be attributed to any individual marker.

Given the redistribution of CD63 at the plasma membrane observed in 231-OE cells (Figure 3), we next investigated whether CD133 expression was associated with changes in CD63^+^ EV populations using complementary approaches. These included single-particle nFCM analysis and MACSPlex detection using a CD63-specific antibody, allowing us to examine the emergence of CD133 /CD63 EVs and assess CD63-associated signals independently of the combined tetraspanin detection used above.

### CD133 expression is associated with the emergence of a CD133 /CD63 EV population

To further characterize changes in EV surface-marker profiles we performed a complementary MACSPlex analysis using an APC-conjugated anti-CD63 antibody for detection, rather than the anti-CD9/CD63/CD81 antibody cocktail used in Figure 4 (Figure 5A, B). EVs were captured using marker-specific antibody-coated beads, and background-corrected APC signals were evaluated relative to the corresponding blank and antibody controls.

**Figure 5.**
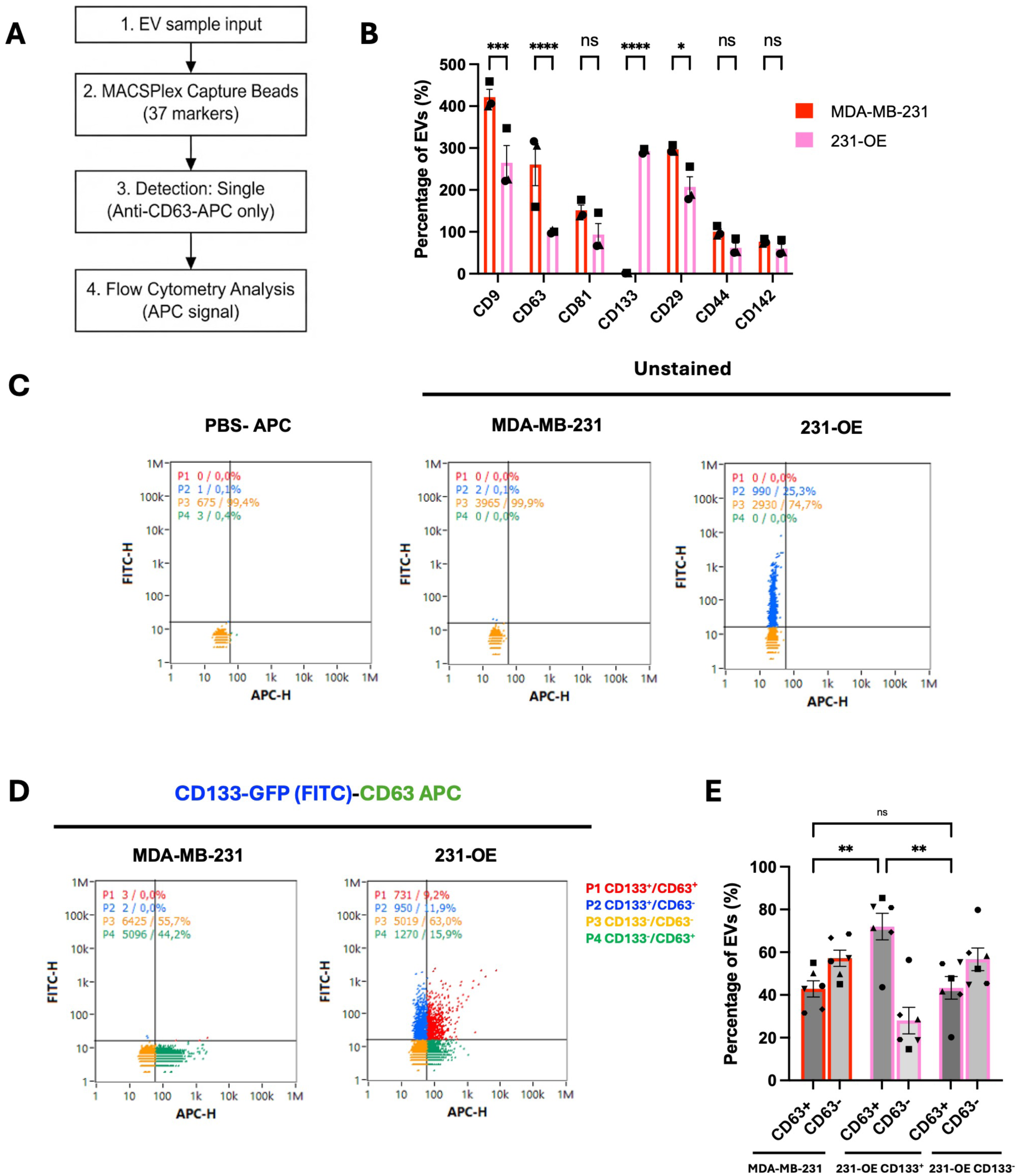
CD133 expression is associated with the emergence of CD133 /CD63 EVs. **A.** Schematic illustrating the MACSPlex workflow. EVs were captured and subsequently detected using APC-labelled anti-CD63 antibody. **B.** MACSPlex analysis detecting tetraspanins, CD133 and other surface markers on EVs from MDA-MB-231 and 231-OE cells. Data are presented as mean ± SEM from 3 independent experiments, analyzed by two-way ANOVA. \*\*\**p* = 0.0001 (CD9); **** *p* < 0.0001 (CD63 and CD133); ns = 0.3798, 0.8188 and 0.9978 (CD81, CD44 and CD142, respectively); \**p* = 0.0027 and 0.0426 (CD29). **C.** Representative flow cytometry controls showing unstained MDA-MB-231 and CD133-GFP-positive 231-OE EVs, used to define the CD133-GFP-positive populations. PBS containing anti-CD63-APC antibody was processed identically to EV-treated samples to assess background fluorescence. **D.** Representative flow cytometry plots from six independent experiments showing CD133-GFP EVs stained with anti-CD63-APC antibody. EV populations were defined as P1 (CD133 /CD63), P2 (CD133 /CD63), P3 (CD133 /CD63), and P4 (CD63 /CD133). **E.** Percentage of CD63 and CD63 EVs within the CD133-positive and CD133-negative EV populations from MDA-MB-231 and 231-OE cells. CD63 EVs were calculated as P1/(P1+P2) and P4/(P3+P4), whereas CD63 EVs corresponded to the complementary fractions. Data are presented as mean ± SEM from six independent experiments. Statistical analysis was performed on CD63 populations only using one-way ANOVA. \*\**p* = 0.0033 and 0.0037 (CD63^+^ MDA-MB-21 vs. CD63^+^ 231-OE CD133^+^ and CD63^+^ 231-OE CD133^+^ vs. CD63^+^ 231-OE CD133); ns = 0.9976 (CD63^+^ MDA-MB-21 vs. CD63^+^ 231-OE CD133).

Comparison of MDA-MB-231 and 231-OE EVs revealed an overall decrease in CD63-detection signals associated with EV populations captured through different surface markers in 231-OE samples compared with parental MDA-MB-231 samples (Figure 5B). Significant reductions were observed for CD9-, CD63- and CD29-capture beads, whereas changes associated with CD81-, CD44- and CD142-capture beads did not reach statistical significance. In contrast, the signal associated with CD133-capture beads increased significantly in 231-OE samples, consistent with the expression of CD133-GFP in these cells. Because the detection antibody specifically recognized CD63, these measurements reflect CD63-detection signals associated with EVs captured through the respective markers and do not directly quantify the abundance of the capture markers themselves. Moreover, this bulk capture-based approach cannot establish whether CD133 and CD63 are present on the same individual EVs.

We therefore examined the co-occurrence of CD133 and CD63 at the single-particle level using nFCM (Figure 5C-E). Because CD133 was expressed as a GFP-tagged protein in the genetically modified cells, GFP fluorescence was used to identify CD133-GFP^+^ EVs, while CD63 was detected using an APC-conjugated antibody, following our recently published protocol for reliable EV labeling (Nevo et al., 2026).

Appropriate controls were used to establish the fluorescence gates and distinguish CD133-GFP-associated events from background signal (Figure 5C). As expected, CD133-GFP-positive events were not detected in parental MDA-MB-231 samples, whereas a distinct CD133-GFP-positive population was detected in 231-OE cells, accounting for approximately 30% of detected events.

Representative nFCM plots distinguished four populations: CD133 /CD63 (P1), CD133 /CD63 (P2), CD133 /CD63 (P3) and CD133 /CD63 (P4) (Figure 5D). Quantification across six independent experiments showed that, in parental MDA-MB-231 samples, CD63 events represented approximately 42% of the total detected EV population. In 231-OE samples, nearly 72% of the CD133 EV fraction was also CD63, whereas approximately 43% of the CD133 EV fraction was CD63 (Figure 5E). The proportion of CD63 events was significantly higher in the CD133 fraction than in both parental MDA-MB-231 samples and the CD133 fraction of 231-OE samples. In contrast, the proportion of CD63 events in the CD133 fraction of 231-OE samples did not differ significantly from that in parental MDA-MB-231 samples. These results indicate that CD63 was more frequently detected in the CD133 than in the CD133 EV fraction, without a detectable increase in the proportion of CD63 events in the CD133 fraction relative to parental cells. Together, these findings highlight distinct CD133 /CD63, CD133 /CD63 and CD133 /CD63 EV populations, underscoring heterogeneity in the distribution of these markers among detected events.

To determine whether this distribution pattern was also observed for another tetraspanin, we performed parallel nFCM analysis using CD9-PE staining (Figure S3). Unlike CD63, CD9 did not show significant differences in the proportion of positive events between the CD133 and CD133 EV fractions in 231-OE samples, or between either fraction and parental MDA-MB-231 samples. Thus, CD9 was distributed more similarly across the CD133-defined EV subpopulations than CD63. This contrast suggests that the preferential association observed between CD133 and CD63 was not mirrored by CD9, arguing against a general enrichment of tetraspanins in CD133 EVs and instead pointing to differences in the distribution of individual EV-associated markers.

Together, these analyses indicate that CD133 expression is associated with changes in of CD63-detection signals in bulk EV preparations and with a distinct distribution of CD63, compared with CD9, among CD133-defined EV subpopulations.

This pattern is consistent with the altered representation of CD63 at protrusive plasma membrane domains observed in 231-OE cells (Figure 3) and raises the possibility that CD133-dependent membrane remodeling contributes to EV heterogeneity. However, these data do not establish a direct mechanistic link between plasma membrane organization and the composition of released EVs.

### CD133 expression is associated with remodeling of the intracellular MVB compartment

Having characterized CD133-associated changes in plasma membrane organization and EV heterogeneity (Figures 3-5), we next investigated whether CD133 expression was also associated with alterations in the intracellular MVB compartment. MVB morphology and abundance were examined by TEM in parental and genetically modified MDA-MB-468 cells (468, 468-KO, and 468-rescue) and MDA-MB-231 cells (231 and 231-OE) (Figure 6A). Quantitative analyses were performed at the single-cell level and included MVB density (number of MVBs per unit cell area), the proportion of cell area occupied by MVBs, individual MVB area, and MVB size distribution (Figure 6B-I).

**Figure 6.**
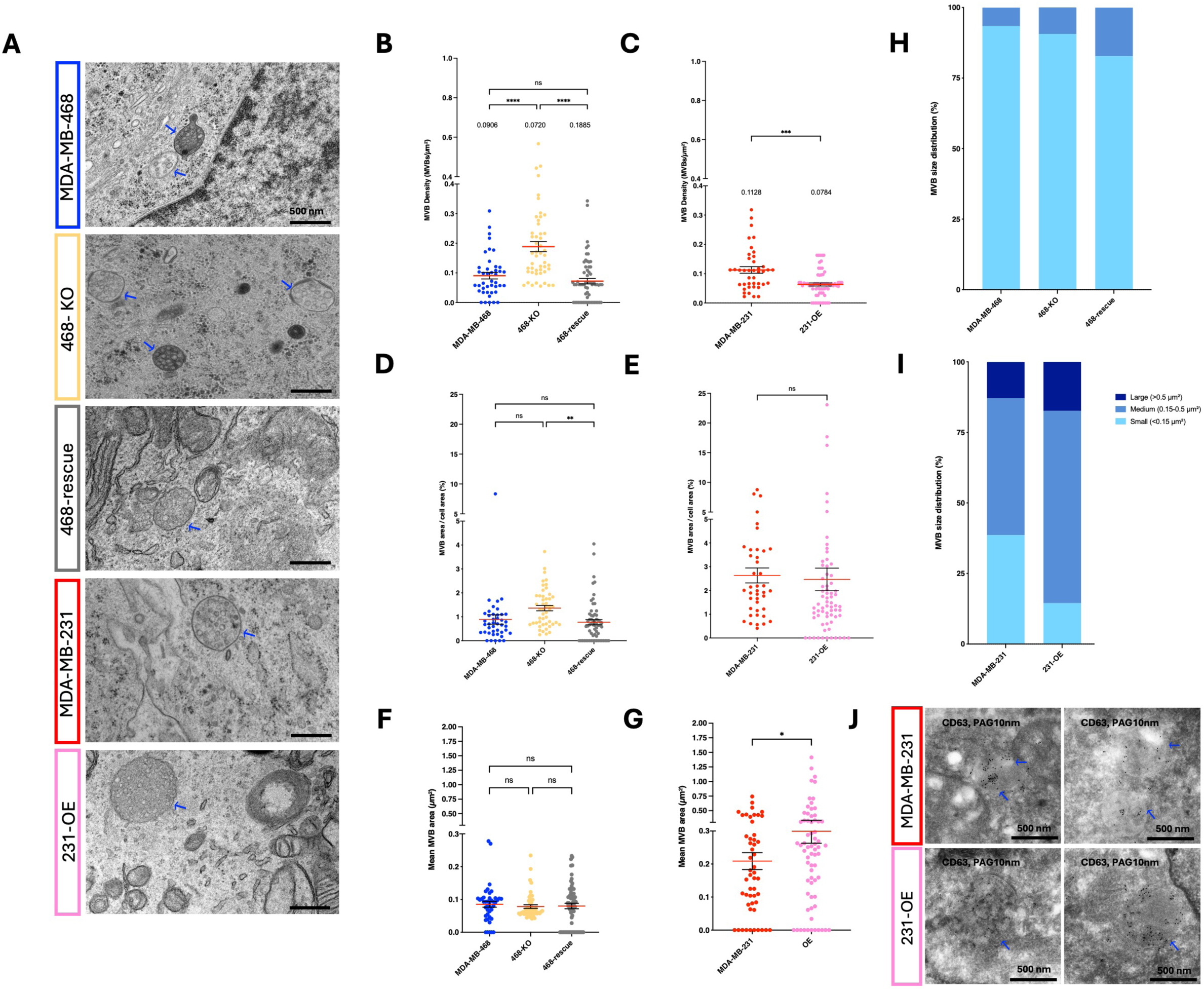
CD133 expression is associated with remodeling of the intracellular MVB compartment. **A.** Representative TEM micrographs of MDA-MB-468, 468-KO, 468-rescue, MDA-MB-231 and 231-OE cells. Blue arrows indicate multivesicular bodies (MVBs). Scale bar: 500 nm. **B-C.** Quantification of MVB density (MVBs/µm²) in (**B**) MDA-MB-468, 468-KO and 468-rescue cells, and (**C**) MDA-MB-231 and 231-OE cells. Data are presented as mean ± SEM from 3 independent experiments, analyzed by one-way ANOVA for MDA-MB-468, 468-KO and 468-rescue cells and by unpaired t-test for MDA-MB-231 and 231-OE cells. \*\*\*\**p* < 0.0001 (MDA-MB-468 *vs*. 468-KO; 468-KO *vs*. 468-rescue); ns = 0.5549 (MDA-MB-468 *vs*. 468-rescue) and \*\*\**p* = 0.0002 (MDA-MB-231 *vs*. 231-OE). **D-E.** Quantification of MVB area relative to cell area (%) in (**D**) MDA-MB-468, 468-KO and rescue cells, and (**E**) MDA-MB-231 and 231-OE cells. Data are presented as mean ± SEM from 3 independent experiments, analyzed by one-way ANOVA for MDA-MB-468, CD133 KO and rescue cells and by unpaired t-test for MDA-MB-231 and 231-OE cells. ns = 0.0545 (MDA-MB-468 *vs*. 468-KO); ns = 0.8249 (MDA-MB-468 *vs*. 468-rescue); \*\**p* = 0.0049 (468-KO *vs*. 468-rescue) and ns = 0.7699 (MDA-MB-231 *vs*. 231-OE). **F-G.** Quantification of MVB area (µm²) in (**F**) MDA-MB-468, 468-KO and 468-rescue cells, and (**G**) MDA-MB-231 and 231-OE cells. Data are presented as mean ± SEM from 3 independent experiments, analyzed by one-way ANOVA for MDA-MB-468, 468-KO and 468-rescue cells and by unpaired t-test for MDA-MB-231 and 231-OE cells. ns = 0.8273 (MDA-MB-468 *vs*. 468-KO); ns = 0.8817 (MDA-MB-468 *vs*. 468-rescue); ns = 0.9880 (468-KO *vs*. 468-rescue) and \**p* = 0.0432 (MDA-MB-231 *vs*. 231-OE). **H-I.** Quantification of MVB size distribution (%) in (H) MDA-MB-468, 468-KO and 468-rescue cells, and (**I**) MDA-MB-231 and 231-OE cells. MVBs were classified by cross-sectional area as small (S, <0.15 µm²), medium (M, 0.15-0.5 µm²) or large (L, >0.5 µm²). **J.** Representative immunogold electron microscopy (IEM) images showing CD63 localization in MVBs of MDA-MB-231 and 231-OE cells.

In the MDA-MB-468 model, CD133 deletion was associated with an increase in MVB density, from approximately 0.09 MVB/µm² in parental cells to 0.19 MVB/µm² in CD133-KO cells, whereas restoration of CD133 expression in 468-rescue cells, reduced MVB density toward the parental level, to approximately 0.07 MVB/µm² in rescue cells (Figure 6B). MVB density was significantly higher in CD133-KO cells than in both parental and rescue cells, whereas the difference between parental and rescue cells was not statistically significant. Similarly, CD133 overexpression in MDA-MB-231 cells was associated with a reduction in MVB density, from approximately 0.11 MVB/µm² in parental cells to 0.06 MVB/µm² in 231-OE cells (Figure 6C).

We next examined the proportion of cell area occupied by MVBs. In the MDA-MB-468 model, this parameter did not differ significantly between parental and 468-KO cells or between parental and 468-rescue cells. However, a significant difference was observed between 468-KO and 468-rescue cells (Figure 6D). In the MDA-MB-231 model, the proportion of cell area occupied by MVBs was comparable between parental and 231-OE cells (Figure 6E). These results indicate that changes in MVB density were not consistently accompanied by corresponding changes in the overall area occupied by MVBs.

We therefore examined individual MVB area. In the MDA-MB-468 model, mean MVB area was approximately 0.08 µm², with no statistically significant differences among parental, 468-KO and 468-rescue cells (Figure 6F). In contrast, MVBs in parental MDA-MB-231 cells had a mean area of approximately 0.21 µm², increasing to approximately 0.30 µm² in 231-OE cells. This increase was statistically significant (Figure 6G). Thus, in MDA-MB-231 cells, CD133 overexpression was associated with both a reduction in MVB density and an increase in mean individual MVB area, whereas no significant change in individual MVB area was detected following CD133 manipulation in the MDA-MB-468 model.

Consistent with the differences in MVB size, analysis of MVB size distributions further highlighted differences between the two cellular models (Figure 6H, I). In MDA-MB-468 model, most MVBs were classified as small, with no large MVBs detected under any of the three conditions (Figure 6H). Specifically, small MVBs accounted for 93.42% of the population in parental cells, 90.63% in 468-KO cells and 82.81% in 468-rescue cells. The corresponding proportions of medium-sized MVBs were 6.58%, 9.38% and 17.19%, respectively.

In contrast, in MDA-MB-231 model, MVB size distribution shifted toward larger structures following CD133 overexpression (Figure 6I). In parental cells, 38.6% of MVBs were small, 48.51% were medium-sized and 12.87% were large. In 231-OE cells, the proportion of small MVBs decreased to 14.49%, while medium-sized and large MVBs accounted for 68.12% and 17.39%, respectively. These observations indicate a shift in the MVB size distribution toward medium-sized structures, accompanied by a more modest increase in the proportion of large MVBs. MVBs were classified according to cross-sectional area as small (<0.15 µm²), medium-sized (0.15–0.5 µm²) or large (>0.5 µm²).

To further characterize MVB morphology, we assessed circularity and aspect ratio. Neither parameter differed significantly across the experimental conditions (Figure S4B–E), indicating that the observed changes in MVB density and size distribution were not accompanied by detectable changes in these measures of MVB shape. The MVB burden index also did not differ significantly between parental and 231-OE cells. In the MDA-MB-468 model, however, the burden index differed significantly between 468-KO and 468-rescue cells, whereas the other reported comparisons were not statistically significant (Figure S4F, G). The proportion of cells containing MVBs was also assessed across the experimental conditions (Figure S4H).

Finally, we examined CD63 labelling in MVBs by immunogold electron microscopy in parental MDA-MB-231 and 231-OE cells (Figure 6J). Quantification showed that the number of CD63-positive gold particles per MVB was significantly lower in 231-OE cells, averaging 14.5 particles per MVB compared with 32.7 in parental cells (Figure S4I). However, when normalized to MVB area, CD63 labelling density was 106.12 gold particles/µm² in parental cells and 95.91 gold particles/µm² in 231-OE cells, with no statistically significant difference between the two conditions (Figure S4J). Thus, the lower number of CD63-positive gold particles per MVB in 231-OE cells was not accompanied by a significant change in CD63 labelling density relative to MVB area.

Together, these analyses show that CD133-associated changes in the intracellular MVB compartment differed between the two TNBC models. In MDA-MB-231 cells, CD133 overexpression was associated with reduced MVB density, increased mean individual MVB area and a shift in size distribution toward medium-sized and, to a lesser extent, large MVBs. In MDA-MB-468 cells, CD133 deletion and rescue were associated with changes in MVB density and size distribution, while mean individual MVB area remained comparable across conditions. The complementary immunogold analysis further revealed a reduction in CD63-positive gold particles per MVB in 231-OE cells, without a significant change in CD63 labelling density per unit MVB area. Collectively, these findings support an association between CD133 expression and remodeling of the intracellular MVB compartment, without establishing a direct mechanistic link between these morphological changes and EV biogenesis or composition.

Taken together, the plasma membrane, EV, and MVB analyses indicate that CD133 expression is associated with coordinated changes in membrane organization, CD63 distribution, EV composition, EV population heterogeneity, and intracellular MVB architecture, supporting a role for CD133 in the regulation of endosomal organization and EV biogenesis.

## DISCUSSION

Extracellular vesicle (EV) heterogeneity is increasingly recognized as an intrinsic feature of EV biology, yet the cellular processes that generate this diversity remain incompletely understood. Although cargo sorting and intracellular trafficking have received considerable attention, EV formation also occurs within a spatially organized membrane system in which protein distribution, membrane curvature and interactions with the cytoskeleton may influence local membrane properties. Our findings support the hypothesis that donor-cell membrane organization contributes to EV heterogeneity. In TNBC cells, CD133 expression was associated with plasma membrane remodeling, a redistribution of CD63 toward membrane protrusions, and distinct distributions of CD133 and CD63 among individual EVs. Changes in MVB organization further indicate that the cellular response to CD133 manipulation extends beyond the plasma membrane, although its nature differs between cellular models. Together, these observations suggest that donor-cell membrane organization may contribute to EV heterogeneity alongside changes in intracellular membrane organization, without establishing a specific EV biogenesis pathway.

The plasma membrane is a spatially organized structure in which lipid composition, protein enrichment, membrane curvature and cytoskeletal interactions can generate domains with distinct properties (Sezgin et al., 2017; van Niel et al., 2018; Rilla, 2021; D’Angelo et al., 2023; Gomez-Duro et al., 2025). Such organization may influence where membrane remodeling occurs and which proteins are locally available during vesicle formation. Consistent with this view, membrane protrusions, including filopodia and microvilli, have been associated with EV release in different cellular contexts (Nishimura et al., 2021; Hurbain et al., 2022). Our results extend this framework by linking CD133 expression to protrusive membrane organization and to the spatial distribution of an EV-associated protein within these structures. The gain- and loss-of-function experiments support a contribution of CD133 to this phenotype, while the incomplete recovery in 468-rescue cells and the differences between the MDA-MB-231 and MDA-MB-468 models indicate that membrane architecture is not determined by CD133 alone.

These findings build on previous work associating CD133 with protrusive membrane structures and cholesterol-rich membrane domains, as well as our demonstration of its contribution to plasma membrane architecture and EV-associated properties in TNBC cells (Gomez-Duro et al., 2026). The present study extends this work by linking CD133-associated membrane remodeling to the spatial redistribution of CD63. In CD133-overexpressing MDA-MB-231 cells, CD63 labeling was disproportionately represented within protrusions relative to the fraction of the plasma membrane occupied by these structures. This suggests that the protrusions differ not only morphologically but also in their molecular organization. Because CD63 is present at both the plasma membrane and intracellular endosomal compartments, including MVBs (Pols and Klumperman, 2009; Mathieu et al., 2021), its distribution may reflect local membrane organization, protein trafficking or retention. The present data do not distinguish between these possibilities, but identify a spatial relationship that may be relevant to EV heterogeneity.

This relationship is particularly relevant to the heterogeneity detected among individual EVs. Single-particle nFCM identified CD133-positive events with and without detectable CD63, as well as CD133-negative/CD63-positive events. CD63 was detected more frequently in the CD133-positive fraction, whereas the proportion of CD63-positive events within the CD133-negative fraction was not significantly different from that in parental-cell preparations. Thus, CD133 expression was associated with a redistribution of CD63 detection across EV subpopulations, rather than a uniform change in CD63 positivity across the entire detected population. This pattern suggests that CD133 positivity identifies a phenotypically distinct subset of detected EVs, rather than a homogeneous population with a uniform marker profile.

The comparison with CD9 further suggests that the distribution of EV-associated proteins is marker-dependent. Unlike CD63, CD9 positivity did not differ significantly across the CD133-defined fractions under the conditions examined. This finding does not establish that CD9 is unaffected by CD133 expression, but it argues against assuming that the distribution of all tetraspanins changes in parallel. More broadly, it highlights the limitations of characterizing EV populations using a single marker or inferring their composition from bulk measurements alone. MACSPlex analysis provided complementary evidence of changes in marker-associated signals, including CD63-detection signal associated with CD133-capture beads. However, this capture-based assay cannot establish that CD133 and CD63 are present on the same individual vesicles. The single-particle analysis provides evidence for their co-detection on a subset of events, while also revealing the diversity of marker combinations within the population.

Importantly, the convergence between plasma membrane imaging and single-particle EV analysis does not establish that CD133+ /CD63+ EVs originate directly from CD63-enriched protrusions. The observed association could arise through local membrane organization, intracellular trafficking, or a combination of these processes. Likewise, the increase in particle output measured following CD133 overexpression does not establish which cellular process accounts for this change or whether the additional particles correspond to a particular EV subpopulation. The relationship between membrane remodeling and EV phenotype therefore remains associative, and direct evidence linking protrusion formation to the production of specific EV populations is still lacking.

The changes in MVB organization provide a complementary perspective on this relationship. CD133 has been detected in endosomal compartments and ILVs, indicating that its cellular distribution is not restricted to the plasma membrane (Karbanova et al., 2025). In our models, CD133 manipulation was associated with changes in MVB density and size distribution, but the responses differed between MDA-MB-231 and MDA-MB-468 cells. These differences suggest that the consequences of CD133 expression for intracellular membrane organization depend on cellular context. They also argue against a uniform effect of CD133 on MVB organization across TNBC models.

The different morphometric parameters provide complementary, rather than interchangeable, descriptions of the MVB compartment. In MDA-MB-231 cells, reduced MVB density and increased individual MVB area were not accompanied by a significant change in the proportion of cell area occupied by MVBs. In the MDA-MB-468 model, differences in MVB density were accompanied by a significant difference in relative MVB area between 468-KO and 468-rescue cells, whereas individual MVB area remained statistically comparable. These findings illustrate that changes in MVB number, size and relative cellular area need not occur in parallel. Although the observed differences are consistent with MVB remodeling, they cannot establish whether MVB formation, maturation, trafficking, fusion or turnover has changed. Nor can static ultrastructural measurements determine whether MVB morphology is functionally linked to the quantity or composition of released EVs. Thus, the MVB data support context-dependent intracellular remodeling but do not identify the mechanism responsible for it or directly explain the EV phenotypes observed.

The CD63 immunogold analysis further illustrates the importance of distinguishing protein labeling per structure from labeling density. Although fewer CD63-positive gold particles were detected per MVB in CD133-overexpressing MDA-MB-231 cells, labeling normalized to MVB area did not differ significantly. The lower number of particles per MVB therefore does not demonstrate reduced CD63 labeling density within the endosomal membrane system or a decrease in total cellular CD63 abundance. Nor can these measurements establish how much CD63 is incorporated into released EVs.

Taken together, the plasma membrane and MVB analyses identify changes in CD63 distribution in distinct cellular compartments, but do not establish that these changes are mechanistically coupled.

Several limitations define the scope of these conclusions. First, the experiments do not directly trace the origin of individual EVs, and therefore cannot establish whether the CD133+/CD63+ population arises from plasma membrane protrusions or intracellular compartments. Second, single-particle measurements depend on labeling specificity, detection sensitivity and gating; the marker-defined populations should consequently be interpreted within the limits of the assay. Third, MACSPlex provides bulk capture-based measurements and cannot resolve marker co-occurrence on individual vesicles. Finally, the different responses of the two TNBC models indicate that the effects associated with CD133 are context-dependent and may involve additional cellular determinants. Future studies combining direct tracking of membrane and endosomal compartments with functional perturbation of membrane organization will be needed to determine how these processes contribute to the formation and release of distinct EV populations.

In conclusion, our study links CD133-associated plasma membrane remodeling to the spatial redistribution of CD63 and to heterogeneity in EV marker combinations, while also identifying context-dependent changes in MVB organization. The preferential detection of CD63 within the CD133-positive EV fraction suggests that CD133-associated changes in the donor-cell membrane system may contribute to differences in EV phenotype. However, the data do not establish a direct connection between the observed membrane protrusions, MVB remodeling and the biogenesis of specific EV populations. Rather than supporting a single route of EV formation, our findings highlight donor-cell membrane organization as a potential contributor to EV heterogeneity and provide a basis for further investigation of the relationship between membrane architecture, intracellular trafficking and EV composition.

## ACKNOWLEDGMENTS

We thank Denis Corbeil, Julien Saint-Pol, Alain Joliot and Guillaume van Niel for helpful discussions. We are also grateful to the CurieCoretech Extracellular Vesicle Platform at Institut Curie for support with the MACSPlex assays, as well as the Flow Cytometry Platform at Institut Curie for technical assistance.

## AUTHOR CONTRIBUTION

M.G.-D., P.B., performed the experiments, P.B, M.G.-D., designed the experiments and analyzed the data. A-S.M. developed the macro used for image analysis. G.D., L.M.-J., C.T., supervised the experiments. G.D. and M.G.-D wrote the paper. G.D. conceived the hypothesis, oversaw the project, supervised all the work and obtained funding. All authors have read the final version of the manuscript and agree to its publication.

## DATA AVAILABILITY STATEMENT

All data generated or analyzed during this study are available in the article and its online supplemental material. Requests for materials should be addressed to G. D’Angelo.

## SUPPLEMENTARY FIGURE LEGEND

**Supplementary Figure 1. CD133 loss reduces plasma membrane protrusion in MDA-MB-468 cells. A**. Representative TEM micrographs of 468-scramble and 468-KO cells. The scramble control corresponds to cells transduced with the non-targeting control CRISPR/Cas9 vector (Gomez-Duro et al., 2025). Blue arrows indicate membrane protrusions. Scale bar: 500 nm. **B.** Quantification of protrusion number on the analyzed membrane and protrusion density (protrusions per µm of membrane) in 468-scramble and 468-KO cells. **C**. Histogram showing the distribution of protrusion length (µm) for scramble and 468-KO cell lines. Data are presented as mean ± SEM from 3 independent experiments, analyzed by one-way ANOVA is shown. \*\*\*\**p* < 0.0001 (MDA-MB-468 *vs*. 468-KO; 468-scramble *vs*. 468-KO) and ns = 0.6451 (MDA-MB-468 *vs*. 468-scramble). **D**. Histogram showing protrusion density (protrusions/µm of membrane) for 468-scramble and 468-KO cell lines. Data are presented as mean ± SEM from 3 independent experiments. Statistical significance was assessed by one-way ANOVA. \*\*\*\**p* < 0.0001 (MDA-MB-468 *vs*. 468-KO), \*\**p* = 0.0034 (468-scramble *vs.* 468-KO) and ns = 0.2112 (MDA-MB-468 *vs.* 468-scramble). **E.** Ratio of protrusive membrane area relative to membrane length (µm) for 468-scramble and 468-KO cell lines. Data are presented as mean ± SEM from 3 independent experiments. Statistical significance was assessed by one-way ANOVA. \*\*\*\**p* < 0.0001 (MDA-MB-468 *vs*. 468-KO; 468-scramble *vs*. 468-KO) and ns = 0.2867 (MDA-MB-468 *vs*. 468-scramble). **F.** Representative ultrathin cryosections of 468-scramble and CD133 KO cells immunolabeled with anti-CD133-PAG (10 nm). Blue arrows indicate CD133 staining localized in membrane protrusions. Right: magnification of the white boxed region. Scale bars are indicated in each image.

**Supplementary Figure 2. Additional IEM images and quantitative measurements underlying the CD63 localization analysis. A.** Representative immunogold electron microscopy (IEM) images showing CD63 localization at the non-protusive (NP) and protrusive (P) plasma membrane in MDA-MB-231 and 231-OE cells. Blue arrowheads indicate CD63-positive gold particles. Right: magnification of the white boxed regions. Scale bars are indicated in each image. **B-C.** Cell-level measurements underlying the CD63 localization analysis in **(B)** MDA-MB-231 cells, and **(C)** in 231-OE cells. The table summarizes the total analyzed membrane length and the corresponding non protrusive (NP) and protrusive (P) regions, including membrane length and CD63-positive gold particles. MDA-MB-231; n = 19; 231-OE; n = 27.

**Supplementary Figure 3. CD133 expression is not associated with a preferential enrichment of CD9 in CD133 EVs. A.** Schematic illustrating the MACSPlex workflow. EVs were captured and subsequently detected using APC-labelled anti-CD9 antibody. **B.** MACSPlex analysis detecting tetraspanins, CD133 and other surface markers on EVs from MDA-MB-231 and 231-OE cells. Data are presented as mean ± SEM from 3 independent experiments, analyzed by two-way ANOVA. ns = >0.9999 (CD9, CD40, CD146 and CD326); ns = 0.9919, 0.9961, 0.9994, 0.9985, 0.9997 (CD63, CD81, CD29, CD44 and CD142 respectively); **** *p* < 0.0001 (CD133). **C.** Representative flow cytometry controls showing PBS containing anti-CD9-PE antibody was processed identically to EV-treated samples to assess background fluorescence. **D.** Representative flow cytometry plots from six independent experiments showing CD133-GFP EVs stained with anti-CD9-PE antibody. EV populations were defined as P1 (CD133 /CD9), P2 (CD133 /CD9), P3 (CD133 /CD9), and P4 (CD9 /CD133). **E.** Percentage of CD9 and CD9 EVs within the CD133 and CD133 EV populations from MDA-MB-231 and 231-OE cells. CD9 EVs were calculated as P1/(P1+P2) and P4/(P3+P4) respectively, whereas CD9 EVs corresponded to the complementary fractions. Data are presented as mean ± SEM from six independent experiments. Statistical analysis was performed on CD9 populations only using one-way ANOVA. ns = 0.9548, 0.1222, 0.1982 (CD9^+^ MDA-MB-21 vs. CD9^+^ 231-OE CD133^+^, CD9^+^ 231-OE CD133^+^ vs. CD9^+^ 231-OE CD133 and CD9^+^ MDA-MB-21 vs. CD9^+^ 231-OE CD133, respectively).

**Supplementary Figure 4. CD133 expression is associated with remodeling of the intracellular MVB compartment. A.** Schematic representation of the MVB circularity scale from 0 to 1. **B-C.** Quantification of MVB circularity in (**B**) MDA-MB-468, 468-KO, and 468-rescue cells, and (**C**) MDA-MB-231 and 231-OE cells. Data are presented as mean ± SEM from 3 independent experiments. Statistical significance was assessed by one-way ANOVA for MDA-MB-468, 468-KO, and 468-rescue cells and by an unpaired t-test for MDA-MB-231 and 231-OE cells. ns = 0.9507 (MDA-MB-468 *vs*. 468-KO); ns = 0.3748 (MDA-MB-468 *vs*. 468-rescue); ns = 0.5024 (468-KO *vs*. 468-rescue) and ns = 0.5166 (MDA-MB-231 *vs*. 231-OE). **D-E.** Quantification of MVB aspect ratio (AR) for MDA-MB-468, 468-KO, and 468-rescue cells, and (**E**) MDA-MB-231 and 231-OE cells. Data are presented as mean ± SEM from 3 independent experiments. Statistical significance was assessed by one-way ANOVA for MDA-MB-468, 468-KO, and 468-rescue cells and by an unpaired t-test for MDA-MB-231 and 231-OE cells. ns = 0.5436 (MDA-MB-468 *vs*. 468-KO); ns = 0.9179 (MDA-MB-468 *vs*. 468-rescue); ns = 0.2728 (468-KO *vs*. 468-rescue) and ns = 0.9152 (MDA-MB-231 *vs*. 231-OE). **F-G.** Quantification of the MVB burden index (a.u.) for (**F**) MDA-MB-468, 468-KO, and 468-rescue cells, and (**G**) MDA-MB-231 and 231-OE cells. Data are presented as mean ± SEM from 3 independent experiments. Statistical significance was assessed by one-way ANOVA for MDA-MB-468, 468-KO, and 468-rescue cells and by an unpaired t-test for MDA-MB-231 and 231-OE cells. ns = 0.0546 (MDA-MB-468 *vs*. 468-KO); ns = 0.8216 (MDA-MB-468 *vs*. 468-rescue); \*\**p* = 0.0048 (468-KO *vs*. 468-rescue) and ns = 0.8344 (MDA-MB-231 *vs*. 231-OE). **H.** Quantification of the percentage of cells containing MVBs, normalized to the analyzed cell surface area, in MDA-MB-468, 468-KO, 468-rescue, MDA-MB-231, and 231-OE cells. **I**. Number of CD63-positive gold-particles within MVBs. Data are presented as mean ± SEM, analyzed by unpaired t-test for MDA-MB-231 and 231-OE cells. \*\*\*\**P* < 0.0001. **J.** Quantification of CD63-positive gold particles within MVBs, expressed as the number of gold particles per µm² of MVB area. Data are presented as mean ± SEM, analyzed by unpaired t-test for MDA-MB-231 and 231-OE cells. ns = 0.4897. Each dot represents one cell (MDA-MB-231 n = 38; 231-OE n = 31).

## REFERENCES

Carney RP, Mizenko RR, Bozkurt BT, Lowe N, Henson T, Arizzi A, Wang A, Tan C, George SC. (2025). Harnessing extracellular vesicle heterogeneity for diagnostic and therapeutic applications. Nat Nanotechnol., Jan;20(1):14–25. doi: 10.1038/s41565-024-01774-3.

Corbeil, D., Marzesco, A.M., Wilsch-Bräuninger, M., & Huttner, W.B. (2010). The intriguing links between prominin-1 (CD133), cholesterol-based membrane microdomains, remodeling of apical plasma membrane protrusions, extracellular membrane particles, and (neuro)epithelial cell differentiation. FEBS Lett., 584(9), 1659–1664. doi: 10.1016/j.febslet.2010.01.050.

D’Angelo, G., Raposo, G., Nishimura, T., & Suetsugu, S. (2023). Protrusion-derived vesicles: new subtype of EVs?. Nat Rev Mol Cell Biol., 24(2), 81–82. doi: 10.1038/s41580-022-00555-x.

D’Angelo, G., Stahl, P.D., & Raposo, G. (2025). The cell biology of extracellular vesicles: A jigsaw puzzle with a myriad of pieces. Curr Opin Cell Biol., 94, 102519. doi: 10.1016/j.ceb.2025.102519.

Dixson AC, Dawson TR, Di Vizio D, Weaver AM. (2023). Context-specific regulation of extracellular vesicle biogenesis and cargo selection. *Nat Rev Mol Cell Biol.*, Jul;24(7):454–476. doi: 10.1038/s41580-023-00576-0.

Fujioka, T., Nishimura, T., Kawana, H. et al. (2025). Efficient cellular transformation via protein delivery through the protrusion-derived extracellular vesicles. Nat Commun., 16, 10900. doi: 10.1038/s41467-025-66351-1.

García-López-de-Haro, C., Fuster-Barceló, C., Rueden, C. T., Heras, J., Ulman, V., & al. (2026). SAMJ: fast image annotation on ImageJ/Fiji via segment anything model. Nat Commun., 17(1), 1–9.

Gomez-Duro, M., Tavares, L. A., Furtado, I. P., Saint-Pol, J., & D’Angelo, G. (2025). Protrusion-derived extracellular vesicles (PD-EVs) and their diverse origins: Key players in cellular communication, cancer progression, and T cell modulation. Biology of the Cell, 117(6), e70018. doi: 19.111/boc.70018.

Gomez-Duro, M., Bergam, P., Sanchez-Redondo, S., Dingli, F., Zhou, A., Joliot, A., Ye, M., Hergueta-Redondo, M., Alves Tavares, L., Loew, D., Dubois, T., Martin-Jaular, L., Peinado, H., Raposo, G., & D’Angelo, G. (2026). CD133 Shapes Extracellular Vesicle Cargo and Angiogenic Function in Basal-Like Triple-Negative Breast Cancer. J Extracell Vesicles, 15(5), e70303. 10.1002/jev2.70303

Hurbain, I., Romão, M., Bergam, P., Heiligenstein, X., & Raposo, G. (2017). Analyzing Lysosome-Related Organelles by Electron Microscopy. Methods in Molecular Biology, 1594, 43–71. doi: 10.1007/978-1-4939-6934-0_4.

Hurbain, I., Macé, A.S., Romao, M., Prince, E., Sengmanivong, L., Ruel, L., … & D’Angelo, G. (2022). Microvilli-derived extracellular vesicles carry Hedgehog morphogenic signals for Drosophila wing imaginal disc development. Curr Biol., 32(2), 361–373. doi: 10.1016/j.cub.2021.11.023. Epub 2021 Dec

Karbanová, J., Thamm, K., Fargeas, C. A., Deniz, I. A., Lorico, A., & Corbeil, D. (2025). Prominosomes-a particular class of extracellular vesicles containing prominin-1/CD133? J Nanobiotechnol., 23(1), 1–39. doi: 10.1186/s12951-025-03102-w.

Marzesco, A.M., Janich, P., Wilsch-Bräuninger, M., Dubreuil, V., Langenfeld, K., Corbeil, D., & Huttner, W.B. (2005). Release of extracellular membrane particles carrying the stem cell marker prominin-1 (CD133) from neural progenitors and other epithelial cells. J Cell Sci., 118(Pt 16), 2849–2858. doi: 10.1242/jcs.02453.

Marzesco, A. M., Wilsch-Bräuninger, M., Dubreuil, V., Janich, P., Langenfeld, K., Thiele, C., Huttner, W. B., & Corbeil, D. (2009). Release of extracellular membrane vesicles from microvilli of epithelial cells is enhanced by depleting membrane cholesterol. FEBS Letters, 583(5), 897–902.

Mathieu M, Névo N, Jouve M, Valenzuela JI, Maurin M, Verweij FJ, Palmulli R, Lankar D, Dingli F, Loew D, Rubinstein E, Boncompain G, Perez F, Théry C. 2021. Specificities of exosome versus small ectosome secretion revealed by live intracellular tracking of CD63 and CD9. Nat Commun., Jul 19;12(1):4389. doi: 10.1038/s41467-021-24384-2.

Nevo N, Zhou A, Ansart N, Cohen-Attali L, Rubinstein E, Guérin C, Martin-Jaular L, Théry C. 2026. Nano-Flow Cytometry of Single Extracellular Vesicles Reveals Subpopulation Differences Across Cell Types and Pharmacological Perturbations. J Extracell Vesicles. Apr;15(4):e70271. doi: 10.1002/jev2.70271.

Nishimura T, Oyama T, Hu HT, Fujioka T, Hanawa-Suetsugu K, Ikeda K, Yamada S, Kawana H, Saigusa D, Ikeda H, Kurata R, Oono-Yakura K, Kitamata M, Kida K, Hikita T, Mizutani K, Yasuhara K, Mimori-Kiyosue Y, Oneyama C, Kurimoto K, Hosokawa Y, Aoki J, Takai Y, Arita M, Suetsugu S. 2021. Filopodium-derived vesicles produced by MIM enhance the migration of recipient cells. Dev Cell. Mar 22;56(6):842-859.e8. doi: 10.1016/j.devcel.2021.02.029.

O’Brien CA, Pollett A, Gallinger S, Dick JE. (2007). A human colon cancer cell capable of initiating tumour growth in immunodeficient mice. Nature. Jan 4;445(7123):106-10. doi: 10.1038/nature05372. Epub 2006 Nov 19.

Pleskač, P., Fargeas, C. A., Veselska, R., Corbeil, D., & Skoda, J. (2024). Emerging roles of prominin-1 (CD133) in the dynamics of plasma membrane architecture and cell signaling pathways in health and disease. Cell Mol Biol Letters, 29(1), 41. doi: 10.1186/s11658-024-00554-0.

Pols MS, Klumperman J. (2009). Trafficking and function of the tetraspanin CD63. Exp Cell Res., May 15;315(9):1584–92. doi: 10.1016/j.yexcr.2008.09.020. Epub 2008 Oct 7. PMID: 18930046.

Reichert, D., J. Scheinpflug, J. Karbanová, D. Freund, M. Bornhäuser, and D. Corbeil. (2016). Tunneling Nanotubes Mediate the Transfer of Stem Cell Marker CD133 Between Hematopoietic Progenitor Cells. Experimental Hematology, 44, no. 11: 1092–1112.e2. doi10.1016/j.exphem.2016.07.006.

Rilla, K. (2021). Diverse plasma membrane protrusions act as platforms for extracellular vesicle shedding. J Extracell Vesicles, 10(11), e12148. doi: 10.1002/jev2.12148.

Sezgin, E., Levental, I., Mayor, S., & Eggeling, C. (2017). The mystery of membrane organization: composition, regulation and roles of lipid rafts. Nat Rev Mol Cell Biol., 18(6), 361–374. doi: 10.1038/nrm.2017.16.

Stringer, C., Wang, T., Michaelos, M. et al. (2021). Cellpose: a generalist algorithm for cellular segmentation. Nat Methods, 18, 100–106. doi: 10.1038/s41492-020-01018-x.

Thamm, K., Šimaitė, D., Karbanová, J., Bermúdez, V., Reichert, D., Morgenstern, A., … & Corbeil, D. (2019). Prominin-1 (CD133) modulates the architecture and dynamics of microvilli. Traffic, 20(1), 39–60. doi: 10.1111/tra.12618. Epub 2018 Nov 14.

van de Wakker, S. I., van Balkom, B. W., de Jager, S. C., Gitz-Francois, J. J., Sluijter, J. P., & Vader, P. (2023). Extracellular vesicle heterogeneity and its impact for regenerative medicine applications. Pharmacological Reviews, 75(6), 1146–1172.

Van Niel, G., D’Angelo, G., & Raposo, G. (2018). Shedding light on the cell biology of extracellular vesicles. Nat Rev Mol Cell Biol., 19(4), 213–228. doi: 10.1038/nrm.2017.125. Epub 2018 Jan 17.

Welsh JA, Goberdhan DCI, O’Driscoll L, et al. (2024). Minimal information for studies of extracellular vesicles (MISEV2023): From basic to advanced approaches. J Extracell Vesicles., 13(2):e12404. doi:10.1002/jev2.12404.

