## Supplementary Figure 1 for "CD133-dependent membrane organization shapes extracellular vesicle heterogeneity in triple-negative breast cancer"

**Supplementary Figure 1. CD133 loss reduces plasma membrane protrusion in MDA-MB-468 cells.**

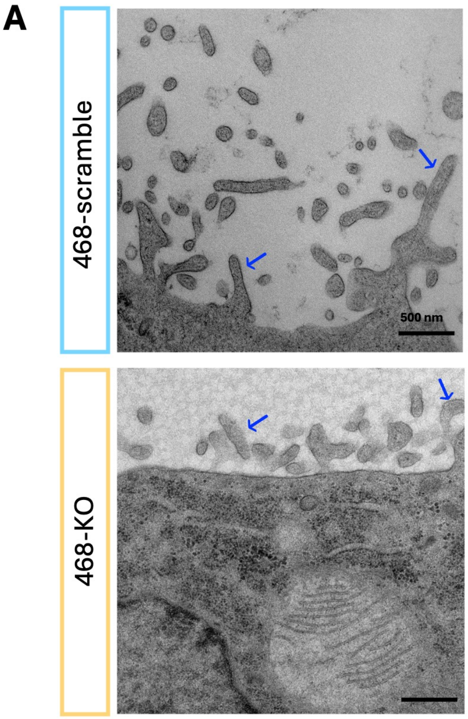


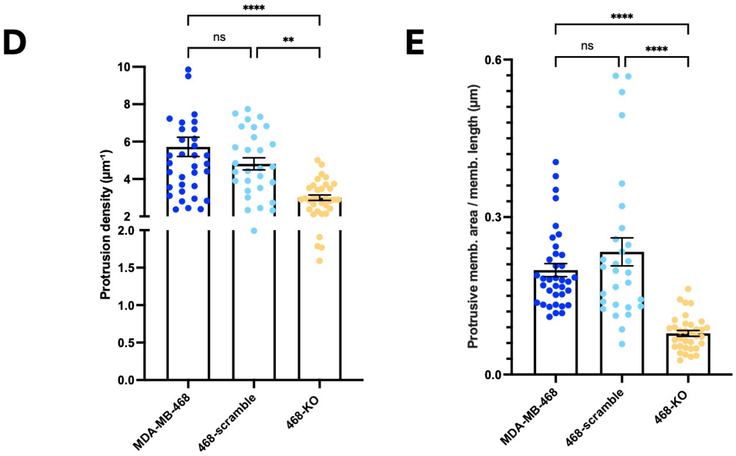


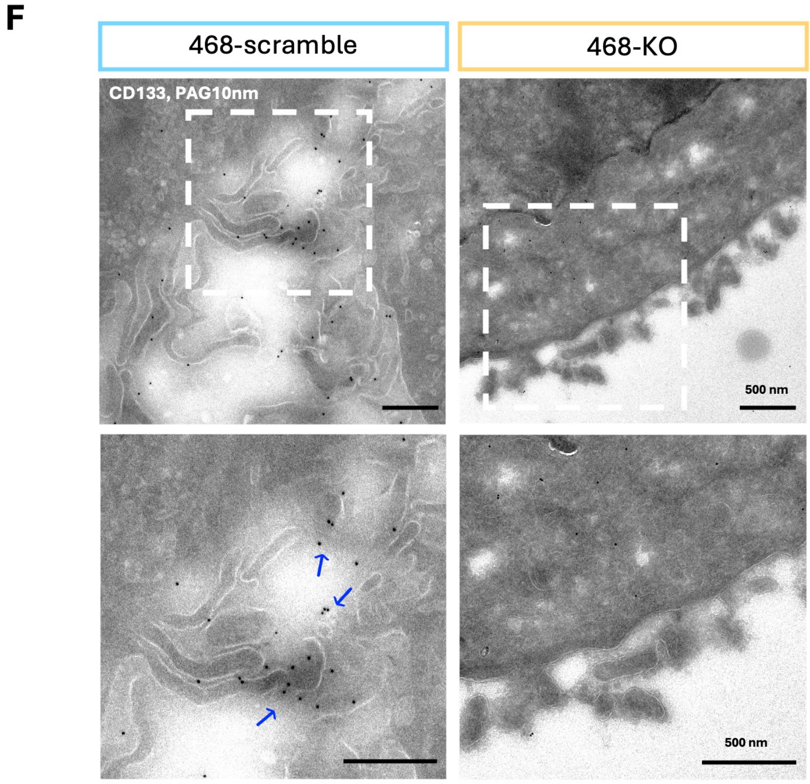


**B**

| Cells | Nb protrus. (analyz. memb. lenght) | Nb protrus. (1 µm) |
|---|---|---|
| 468-scramble | 813 | 4.6 |
| 468-KO | 515 | 2.9 |

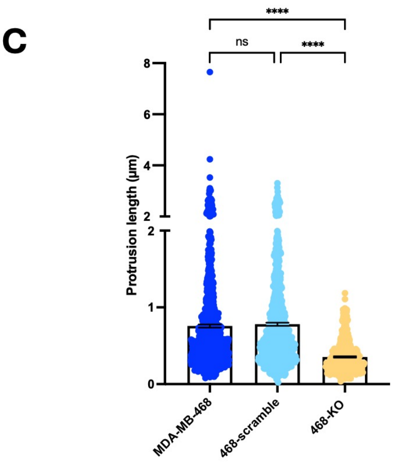
