## Supplementary Figure 2 for "CD133-dependent membrane organization shapes extracellular vesicle heterogeneity in triple-negative breast cancer"

**Supplementary Figure 2. Additional IEM images and quantitative measurements underlying the CD63 localization analysis**

**A**

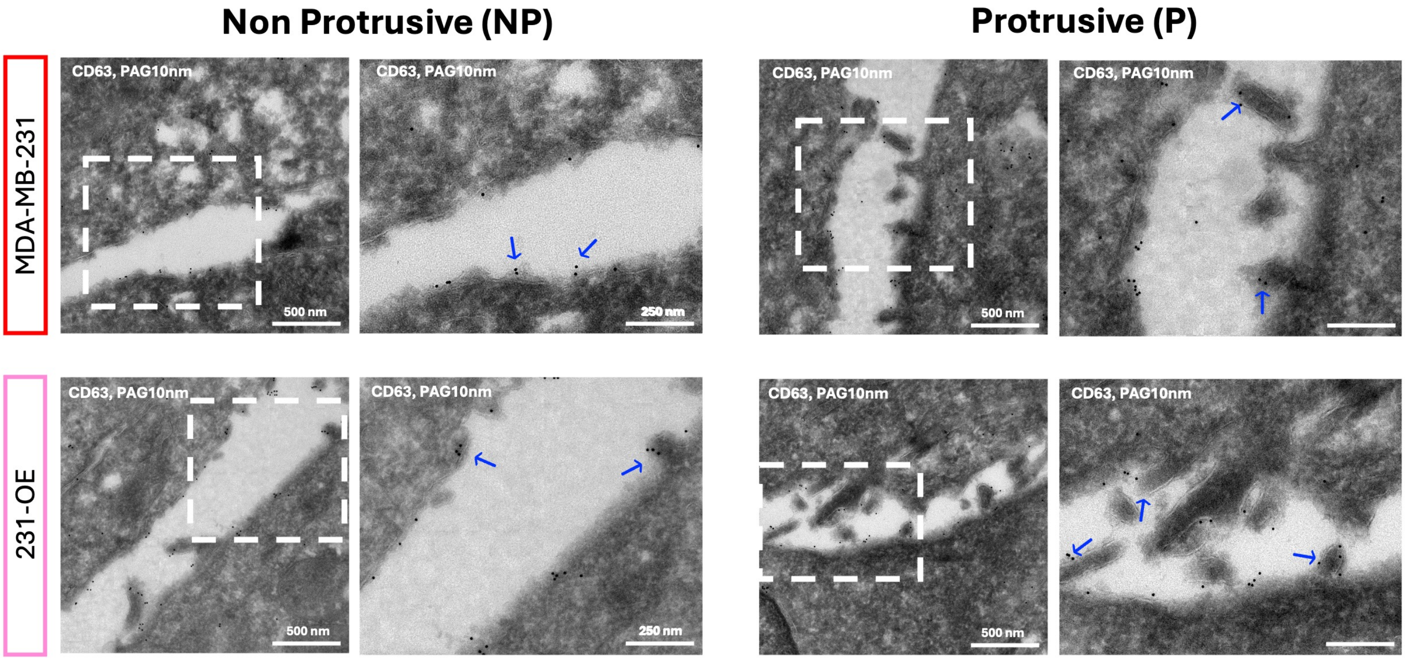


**B**

MDA-MB-231 n= 19

| Total Membrane Length | Membrane Length | | CD63 beads | |
|---|---|---|---|---|
| | NP | P | NP | P |
| 6,7 | 6,7 | 0,0 | 40,0 | 0,0 |
| 5,0 | 5,0 | 0,0 | 23,0 | 0,0 |
| 6,2 | 5,0 | 1,2 | 7,0 | 5,0 |
| 2,0 | 2,0 | 0,0 | 33,0 | 0,0 |
| 4,6 | 4,0 | 0,6 | 27,0 | 2,0 |
| 2,6 | 2,6 | 0,0 | 49,0 | 0,0 |
| 4,0 | 4,0 | 0,0 | 23,0 | 0,0 |
| 4,9 | 4,9 | 0,0 | 71,0 | 0,0 |
| 7,1 | 6,2 | 0,9 | 34,0 | 4,0 |
| 4,5 | 4,5 | 0,0 | 32,0 | 0,0 |
| 5,6 | 5,0 | 0,6 | 70,0 | 2,0 |
| 7,6 | 6,6 | 1,0 | 36,0 | 4,0 |
| 7,4 | 5,9 | 1,5 | 26,0 | 7,0 |
| 2,3 | 2,3 | 0,0 | 45,0 | 0,0 |
| 4,0 | 4,0 | 0,0 | 65,0 | 0,0 |
| 2,9 | 2,9 | 0,0 | 92,0 | 0,0 |
| 5,0 | 5,0 | 0,0 | 71,0 | 0,0 |
| 5,8 | 4,9 | 0,9 | 83,0 | 7,0 |
| 2,2 | 2,2 | 0,0 | 34,0 | 0,0 |

**C**

231-OE n= 27

| Total Membrane Length | Membrane Length | | CD63 beads | |
|---|---|---|---|---|
| | NP | P | NP | P |
| 10,3 | 6,8 | 3,4 | 19,0 | 26,0 |
| 10,7 | 7,3 | 3,4 | 24,0 | 40,0 |
| 5,0 | 4,1 | 0,9 | 8,0 | 6,0 |
| 13,1 | 7,3 | 5,9 | 17,0 | 30,0 |
| 13,4 | 6,9 | 6,4 | 14,0 | 24,0 |
| 10,9 | 6,8 | 4,1 | 27,0 | 20,0 |
| 3,9 | 3,9 | 0,0 | 4,0 | 0,0 |
| 2,1 | 1,6 | 0,5 | 3,0 | 3,0 |
| 4,7 | 3,9 | 0,8 | 4,0 | 3,0 |
| 7,6 | 5,3 | 2,3 | 21,0 | 42,0 |
| 4,8 | 3,9 | 0,9 | 27,0 | 11,0 |
| 13,7 | 4,7 | 8,9 | 12,0 | 54,0 |
| 5,6 | 4,7 | 0,9 | 17,0 | 7,0 |
| 11,8 | 2,7 | 9,1 | 13,0 | 117,0 |
| 20,5 | 5,0 | 15,5 | 10,0 | 136,0 |
| 4,3 | 3,8 | 0,5 | 20,0 | 2,0 |
| 6,8 | 5,7 | 1,1 | 20,0 | 21,0 |
| 7,7 | 7,0 | 0,7 | 12,0 | 2,0 |
| 11,3 | 4,7 | 6,6 | 21,0 | 52,0 |
| 3,2 | 3,2 | 0,0 | 22,0 | 0,0 |
| 3,8 | 3,8 | 0,0 | 7,0 | 0,0 |
| 7,1 | 5,4 | 1,6 | 12,0 | 7,0 |
| 9,7 | 5,3 | 4,4 | 14,0 | 30,0 |
| 8,3 | 6,2 | 2,1 | 6,0 | 12,0 |
| 2,3 | 2,3 | 0,0 | 0,0 | 0,0 |
| 11,0 | 7,1 | 3,9 | 40,0 | 11,0 |
| 5,1 | 3,9 | 1,2 | 13,0 | 17,0 |
