## Supplementary Figure 3 for "CD133-dependent membrane organization shapes extracellular vesicle heterogeneity in triple-negative breast cancer"

**Supplementary Figure 3. CD133 expression is not associated with a preferential enrichment of CD9 in CD133⁺ EVs.**

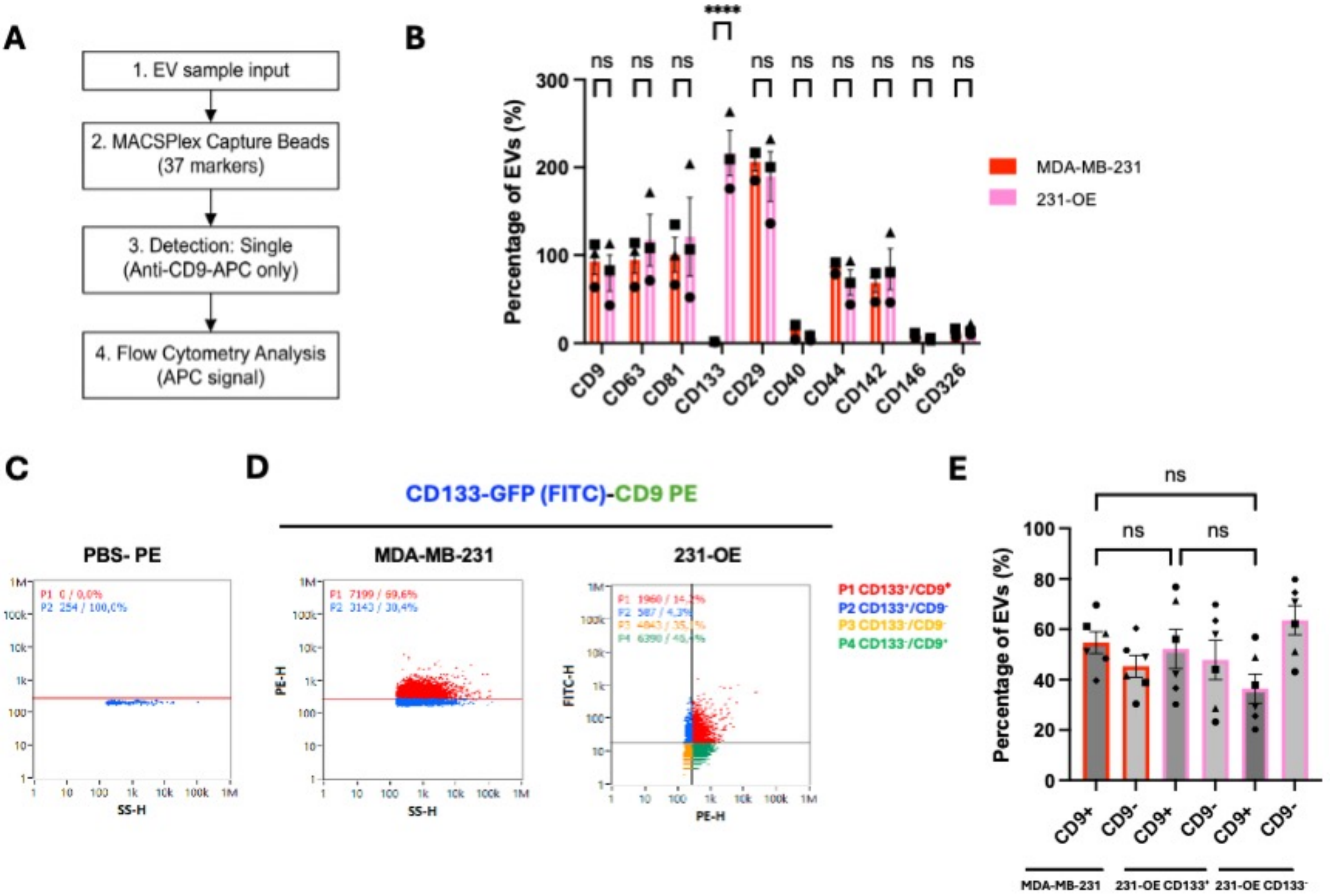
