## Supplementary Figure 4 for "CD133-dependent membrane organization shapes extracellular vesicle heterogeneity in triple-negative breast cancer"

### Supplementary Figure 4. CD133 expression is associated with remodeling of the intracellular MVB compartment

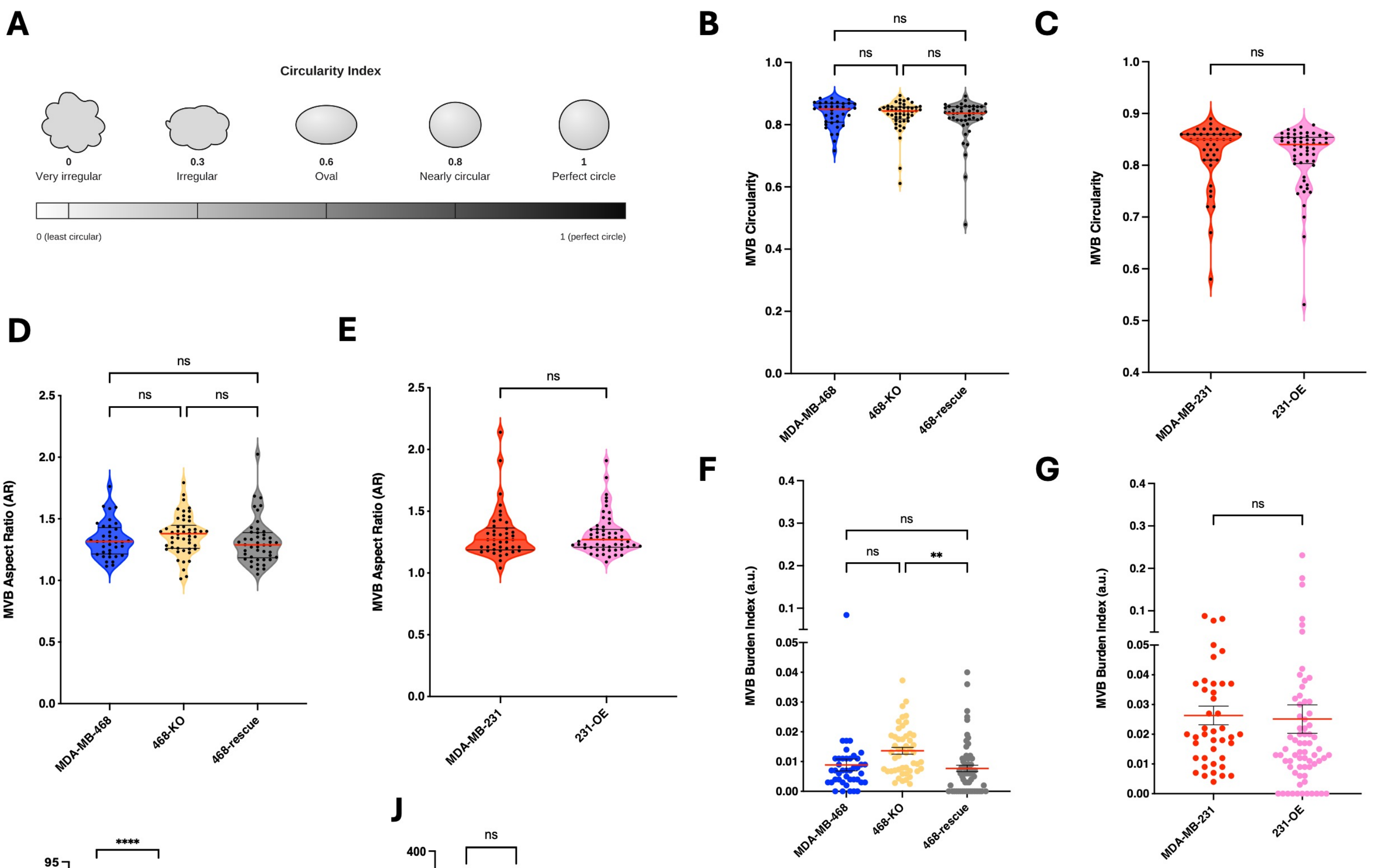


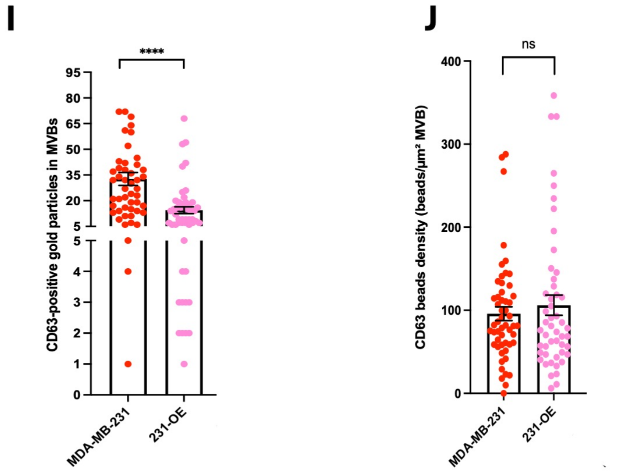


H

| Cell line | % Cells with MVB |
|---|---|
| MDA-MB-468 | 88,10 |
| 468-KO | 100 |
| 468-rescue | 71,43 |
| MDA-MB-231 | 100 |
| 231-OE | 83,82 |
